# Nuclear Factor I genes drive chondrogenic cell-fate commitment

**DOI:** 10.64898/2026.04.21.719911

**Authors:** Rick Mulders, Mathew Nickel-Maunu, Marcella van Hoolwerff, Giorgia Mazzini, Lucas Klomp, Hil Meijer, Janine Post, Yolande F. M. Ramos, Ingrid Meulenbelt

**Affiliations:** Dept. of Biomedical Data Sciences, Section Molecular Epidemiology, Leiden University Medical Center, Leiden, The Netherlands

## Abstract

Human induced pluripotent stem cells (hiPSCs) offer a powerful platform to model chondrogenesis and enable regenerative strategies, yet regulation of cell-fate commitment remains elusive. Here, we systematically mapped cell-fate trajectories from 7 time points during a 49-day chondrogenic hiPSC differentiation protocol using single-nucleus multimodal transcriptomic and chromatin accessibility profiling (scRNA-seq and scATAC-seq). Integrative analysis of dynamics revealed branching differentiation trajectories with clear bifurcation points and distinct cell-fates. Notably, the chondrogenic trajectory originated at day 6 as a neurogenic development and branched off at day 21 to a chondrogenic cell-fate. Through transcription factor activity analysis (TFAA) and cis-co-accessibility networks, we found that *NFIA* and *NFIB* drove chondrogenic distinction, exhibited in both modalities as directly targeting chondrogenic genes such as *COMP*, *FIBIN*, *VIM.* This was then confirmed by experimental validation as modulation of *NFIA* expression at this point further enhanced chondrocyte formation. Together, our study provides a high-resolution multimodal atlas of chondrogenic differentiation and identified critical transcriptional regulators that can now be leveraged to implement regenerative cartilage therapies from hiPSCs.

## Introduction

Osteoarthritis is a painful and debilitating joint disease. Articular cartilage of the covering long bone joint facing surfaces is progressively worn down until bare bone is exposed. Currently, no effective treatment exists, with total joint replacement surgery as the only option at the end stage of disease progression. Much effort has been made to produce regenerative therapies for affected joints, replenishing worn cartilage with new material. These approaches use autologous articular chondrocytes or mesenchymal stem cells (MSCs) as cell source^1,2^. While initial results seem promising, scalability is a problem. Moreover, chondrocytes have a tendency to dedifferentiate in culture^3^ and MSCs are prone to produce hypertrophic chondrocytes.^4^ Therefore, a stable source of off-the-shelf high quality articular cartilage is a much desired development. In previous work we have produced high quality neo-cartilage from human induced pluripotent stem cells (hiPSCs)^5,6^, achieving an 85% methylome similarity to primary chondrocytes. The 49-day protocol is derived from Adkar *et al*. 2019^7^ and drives hiPSCs, via the paraxial mesoderm, early somite and sclerotome, to chondroprogenitors (hiCPCs) in monolayer cultures up until day-14. Thereafter, cell aggregates are picked for maturation in suspension towards hiPSC-derived chondrocytes (hiCHOs).^5,7^ Despite the high similarity between induced chondrocytes and primary chondrocytes, some remaining discrepancy was highlighted by increased expression levels of neuronal markers like *KIF1A* and *NRXN1.*^5^. This means that at the end of the protocol, at day 49, there could be either a heterogenous population of cells consisting e.g. of ‘on-target’ mature chondrocytes and ‘off-target’ cells or a homogeneous population of cells with a more stem-like or immature cellular phenotype. Such uncertainty in cell population composition prevents application of hiPSC-derived tissues for regenerative solutions in a clinical setting.

To determine and understand key modifiable factors that direct and maintain hiPS-cell differentiation into stable chondrogenic cell fates, we applied RNA-sequencing and assay for transposase-accessible chromatin (ATAC)-sequencing simultaneously in single nuclei at 7 timepoints throughout the 49-day-differentiation. By exploiting this dataset, we mapped the cell-fate trajectories that arose during the differentiation process and identified the dynamic changes in expression that marked cell-fate commitments. Then, by taking advantage of the temporal and multimodality aspects of our single-cell data we captured and experimentally validated critical drivers of chondrogenic cell-fate commitment. Together our study provides critical modifiable transcriptional regulators that can now be leveraged to implement regenerative cartilage therapies from hiPSCs.

## Results

### Single-nucleus multimodal sequencing of chondrogenic differentiation

To determine and understand key modifiable factors that direct hiPSC chondrogenic differentiation, we generated a single-nucleus multimodal dataset from samples taken over the course of the entire 49-day-differentiation protocol. ^9,10^ As shown in **Fig. 1A**, rapid media changes in the first 3 days of chondrogenic differentiation ensure developmental progression to the early somite stage, followed by media changes that were spaced out further to facilitate gradual differentiation to what was presumed to be sclerotome (day-6) and chondroprogenitors (hiCPCs, day-14), all in 2D cultures. Thereafter, hiCPC aggregates were put into suspension cultures for 3D maturation to chondrocytes (hiCHOs), up until day-49. To allow simultaneous nuclei isolation, library preparation, and sequencing procedures of cells from days 0, 3, 6, 8, 14, 21, and 49, we staggered differentiation so that a sample of each day was ready to harvest at the same time. As shown in **Fig. 1B**, after 49 days of hiPSC chondrogenic differentiation Alcian Blue-positive neo-cartilage organoids were formed, albeit that still non-cartilage cell aggregates were present in variable quantities.

**Figure 1:**
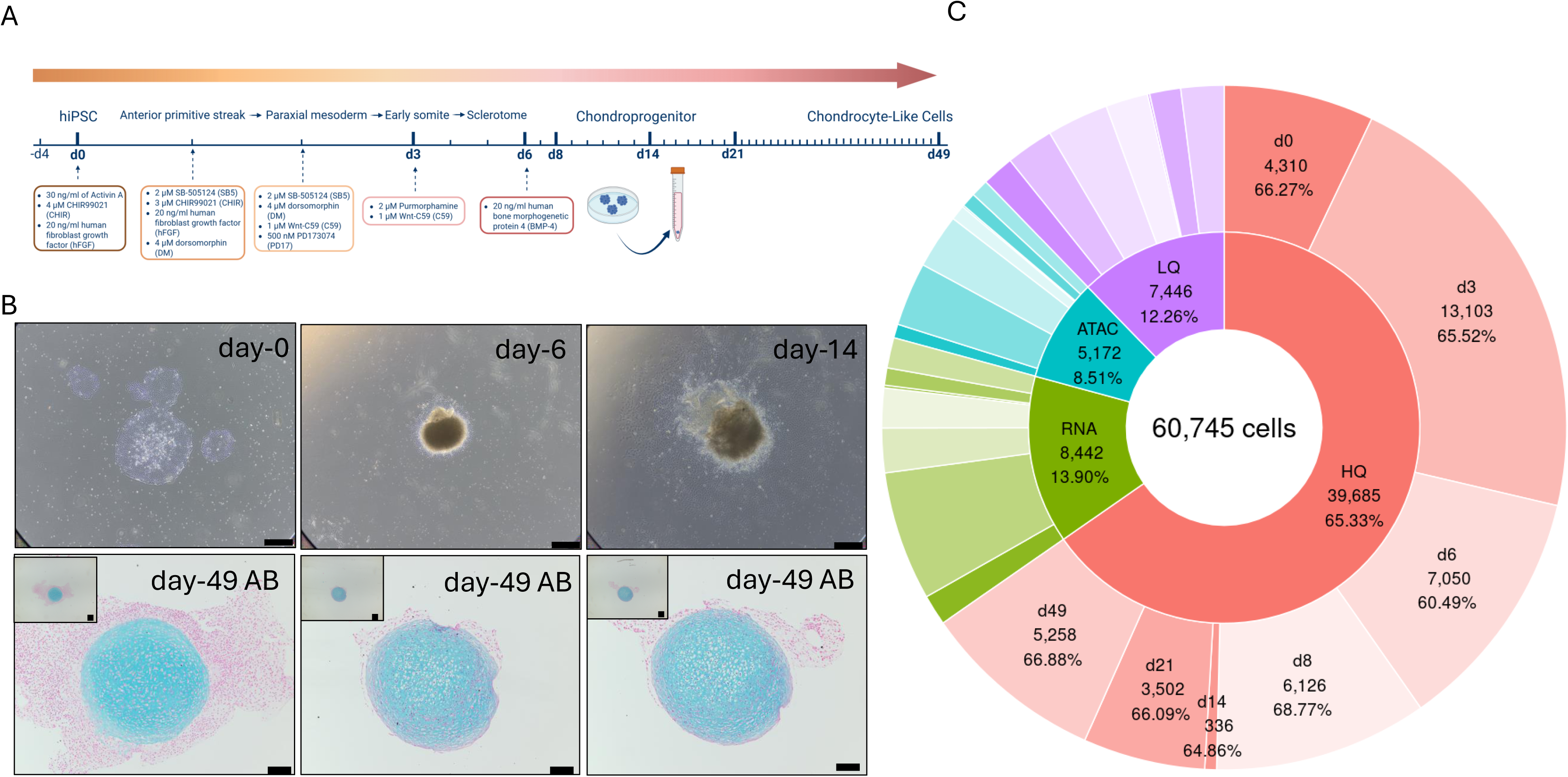
Generating a high-quality dataset of chondrogenic differentiation *in vitro*. **A**) Diagram of chondrogenic differentiation of hiPSCs for 49 days. Included are the compounds added to the differentiation medium at each point. At day 14 cell aggregates from 2D culturing conditions are cultured in 3D for the remainder of the differentiation. Indicated above the timeline are the cell types expected at that stage of differentiation. The numbered ticks represent timepoints of harvesting. **B**) Histological images of cells during differentiation. The top row shows cell colonies at day 0 (left), day 6 (middle), and day 14 (right), black bars are 500µm in length. The bottom row show images of Alcian Blue (AB) staining of neocartilages at 49 days of differentiation, black bars are 100µm in length. **C**) Donut plot with two rings showing the number of cells that passed quality control. The inner ring is for the sum of all timepoints and is sectioned as high quality in both modalities (HQ, red), only high-quality RNA-seq data (RNA, green), only high-quality in ATAC-seq data (ATAC, blue), and low quality in both modalities (LQ, purple). The outer ring shows the same categories; HQ, RNA, ATAC, LQ in corresponding colours to the inner ring, sections are each timepoint in chronological order, clockwise; day-0, day-3, day-6, day-8, day-14, day-21, day-49.

To be able to capture cell-fate decisions during hiPSC chondrogenic differentiation, 10,000 cells were submitted for library preparation and single nucleus multimodal sequencing at day-0, 3, 6, 8, 14, 21, and 49. In retrospect, the number of nuclei sequenced generally ranged between 5,299 (day-21) and 11,654 (day-6) cells, with the exception of 20,000 cells at day-3, the maximum detectable capacity, and 518 cells at day-14. As shown in **Fig. 1C** and **Supplementary Figure 1,** of the 60,745 cells that were sequenced in total, 39,685 cells passed quality control in both modalities making up 65% of cells both by numbers and mean (σ: 2.6%). Between modalities there was an 85% (σ: 2.6%) consensus in high quality cells. Every timepoint had a similar percentage of high-quality cells (60-68%), despite fluctuating numbers of total cell count.

### Continual mapping of hiPSC chondrogenic differentiation

To map chondrogenic differentiation process we performed clustering analysis on all high quality cells together. Herein, dimensionality reduction for the mRNA and ATAC modalities were first analysed separately, using uniform manifold approximation and projection (UMAP). As shown in **Fig. 2A** and **Fig. 2B**, the mRNA and ATAC projections are similar in that they each show a chronological progression of timepoints by adjacency yet still display unique features per modality, particularly at a local scale. Next, we leveraged the advantage of simultaneous multimodal sequencing to forgo integration across the mRNA and ATAC modalities and use weighted nearest-neighbour networks (WNN) to let both modalities inform the UMAP across the 49-day differentiation. In this low dimensional cell projection (**Fig. 2C**) each timepoint appeared in chronological order. While days 0 and 3 were disconnected from each other and the proceeding timepoints, from day-6 onwards all timepoints were connected, as such capturing the full extent of transcriptomic and epigenetic change which occurs during differentiation. Herein, day-6 and day-8 mixed more than other timepoints, separating into two visually distinct clusters of mixed origin. This structure was persistent across the single modality and WNN projections. To subsequently cluster cells by similarity of the molecular landscape regardless of the timepoint of origin, we used Louvain clustering. As shown in **Fig. 2D**, the Louvain clustering resulted in 13 clusters numbered from 0-12. Although these clusters matched the visual clusters of the UMAP projection closely, some timepoints were separated into several clusters e.g. day-49 was separated in cluster-9, cluster-8, and cluster-5 whereas other timepoints were conjoined into specific clusters, e.g. cluster-6 consisted of cells from day-6, day-8, and day-14.

**Figure 2:**
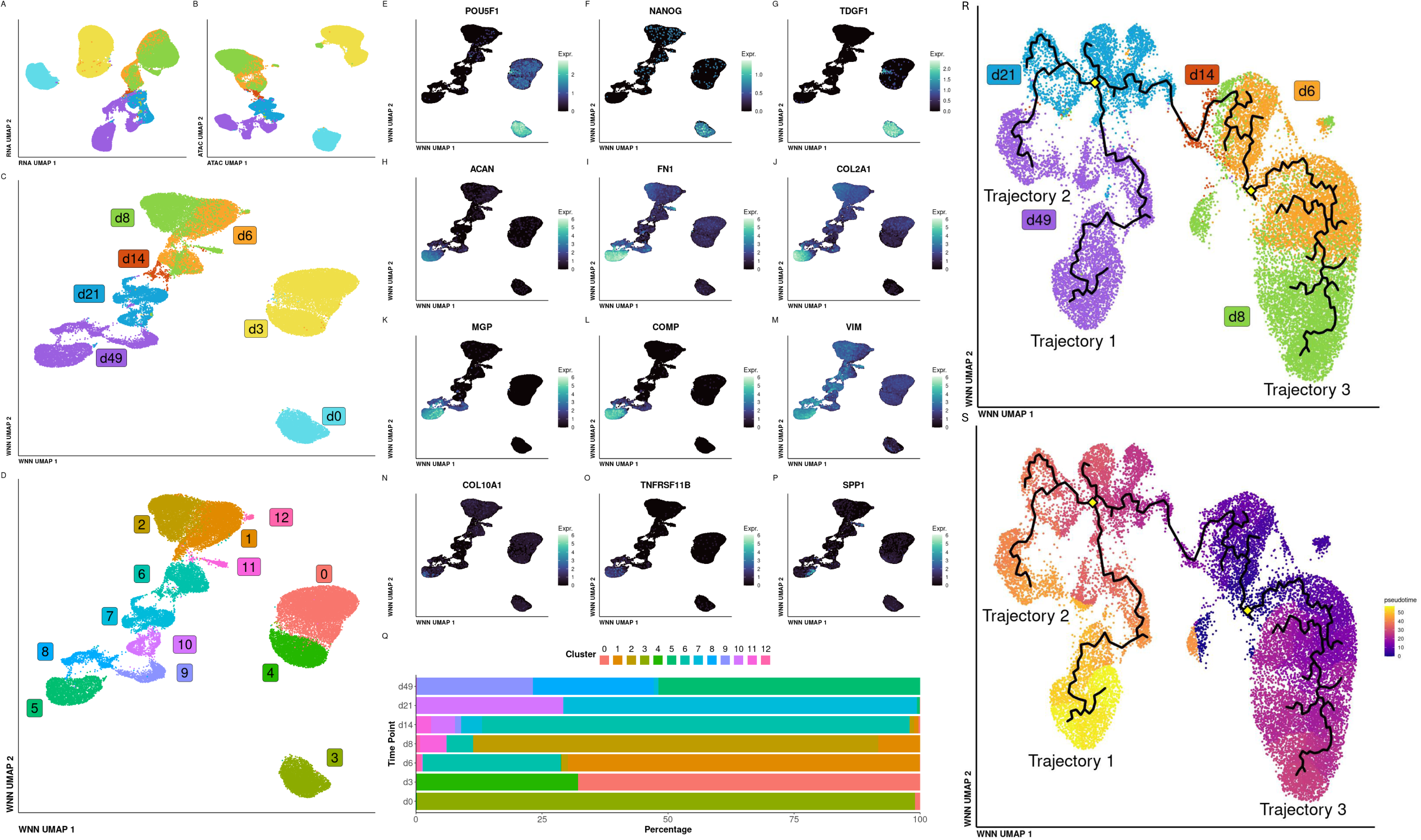
Exploration of single-nucleus multimodal sequencing data of all timepoints; day-0, day-3, day-6, day-8, day-14, day-21, day-49. **A**) UMAP projection of RNA sequencing data, colours align with the labeling in 2C. **B**) UMAP projection of ATAC sequencing data, colours align with labelling in 2C. **C**) UMAP projection of RNA and ATAC data combined by WNN, labels correspond with cell colour in figure 2A, 2B, and 2C. **D**) WNN UMAP projection showing louvain (res 0.2) clusters, labels correspond with cell colour. **E-G**) WNN UMAP projections showing SCT expression for specific genes; *POU5F1* (E), *NANOG* (F), *TDGF1* (G). **H-P**) WNN UMAP projections showing SCT expression for specific genes, the colour gradient is scaled to the maximum expression in this whole set of genes; *ACAN* (H), *FN1* (I), *COL2A1* (J), *MGP* (K), *COMP* (L), *VIM* (M), *COL6A1* (N), *TNFRSF11B* (O), *SPP1* (P) **Q**) WNN UMAP projection showing louvain (res 0.2) clusters, labels correspond with cell colour. **R**) WNN UMAP projection of expression and open chromatin data of timepoints day-6, day-8, day-14, day-21, and day-49 with inferred trajectory coloured by timepoint. Label colour corresponds to cell colour. Branching trajectory in black lines, yellow diamonds indicate two major points of bifurcation, trajectory labels in open space indicating closest label. **S**) WNN UMAP projection of timepoints day-6, day-8, day-14, day-21, and day-49 with inferred trajectory coloured by pseudotime.

To capture cellular characteristics between day-1 and day-49 during hiPSC chondrogenic differentiation, we plotted the top ten differential expressed coding genes per cell cluster (**Supplementary Figure 2**). At the start of differentiation we showed that cluster-3 cells were primarily populated by day-0 cells i.e. hiPSCs, as marked by pluripotency markers *POU5F1* (**Fig. 2E**), *NANOG* (**Fig. 2F**), and *TDGF1* (**Fig. 2G**). On the other hand, at the end of differentiation, we showed that clusters-5, cluster-8, and cluster-9 were primarily populated by day-49 cells i.e. hiCHOs as specifically marked by *ACAN* (**Fig. 2H**), *FN1* (**Fig. 2I**), *COL2A1* (**Fig. 2J**), *MGP* (**Fig. 2K**), and *COMP* (**Fig. 2L**). Notably, for *FN1*, and *COL2A1* a secondary region of moderate expression at day-8 was visible whereas *VIM* (**Fig. 2M**) followed a continually increasing expression pattern from day-6 onwards, while maxing out at day-49 at the terminus of the differentiation. On a different note, hypertrophy markers such as *COL10A1* (**Fig. 2N**), *TNFRSF11B* (**Fig. 2O**), and *SPP1* (**Fig. 2P**) were lowly expressed overall. Important to note is also that pluripotent cells from day-0 and/or day-3 only populated the early clusters (cluster-3, cluster-4, and cluster-0) and were not present in day-49 chondrogenic clusters (cluster-9, cluster-8, and cluster-5) (**Fig. 2Q**).

To trace trajectories of differentiation that link terminal cell clusters to intermediary ones as far back as continuity by similarity allowed, we performed trajectory inference. Due to the gaps between day-0 and day-3 with the other timepoints, we focused on timepoints day-6, 8, 14, 21, and 49. As shown in **Fig. 2R**, trajectory inference resulted in a branched trajectory connecting each of the timepoints with two major points of bifurcation (yellow diamonds) resulting in three trajectories. As shown in **Fig. 2S**, two of these trajectories started simultaneously at day-6 and progressed beyond day-14 where they encounter a bifurcation at day-21. Thereafter we defined **Trajectory-1** as the branch that terminated with a specific cell-fate at pseudo-time unit 56 (yellow) and **Trajectory-2** as the branch terminated with a specific cell-fate at pseudo-time unit 45 (orange). The fact that Trajectory-1 and Trajectory-2 had a difference in pseudo-time but occurred simultaneously in real time (day-49) suggested a difference in differentiation rate. Finally, we observed a third trajectory (**Trajectory-3**) that started at the day-6 bifurcation, progressed to day-8 but then terminated to a specific cell-fate at pseudo-time unit 28 (pink) that was not found back in day-14 cells (**Fig. 2S**).

### Chondrogenic cell-fate commitment emerges from a neurogenic development

To characterize the dynamic changes in gene expression along the three trajectories in a data-driven manner, we generated gene co-expression modules by clustering across all cells with highly autocorrelated genes using Moran’s I (I > 0.25, FDR < 0.001). Subsequently, gene enrichment analysis (Gene ontology: biological process, GO:BP) was performed on co-expression modules to identify associated pathways. As shown in **Fig. 3A**, four significant gene co-expression modules were found (**Supplementary Table 1**). Among the top most representative genes of the prominent **Module-1** genes (red dots, 308 genes) we recognized chondrogenic markers such as *FN1, COL2A1, SOX9, VIM, MGP*, and *COMP*. Gene set enrichment analyses (**Fig. 3B**) confirmed these genes to act in extracellular matrix organisation (GO:0030198), cartilage development (GO:0051216) and ossification (GO:0001503; **Supplementary Table 2**). Among the **Module-2** genes (green dots, 233 genes) we recognized neurogenic genes such *NRXN1, STMN2*, *LRP2,* and *KIF1A* with enrichment (**Fig. 3C**) for basal regulation of neuron projection development (GO:0010975) and CNS neuron differentiation (GO:0021953) and more advanced neurogenic development such as axonogenesis (GO:0007409) and synapse organization (GO:0050808) as most significant results (**Supplementary Table 3**). Among the **Module-3** genes (blue dots, 164 genes) we recognized mesenchymal developmental genes such as *TWIST1*, *YAP1* and *MEOX1*, but also neural crest related genes such as *MDK* and *PTRG*. Subsequent pathway analyses (**Fig. 3D**) confirmed significant enrichment for genes involved in Neural Crest Cell differentiation (GO:0014033) and Epithelial to Mesenchymal transition (GO:0001837). **Module-4** (purple dots, 70 genes), was defined by ribosomal transcripts which can indicate a high degree of translation and proliferation further confirmed by gene enrichment for cytoplasmic translation (GO:0002181), signal transduction by p53 class mediator (GO:0072331), and regulation of amide metabolic process (GO:0034248; **Fig. 3E** and **Supplementary Table 5**).

**Figure 3:**
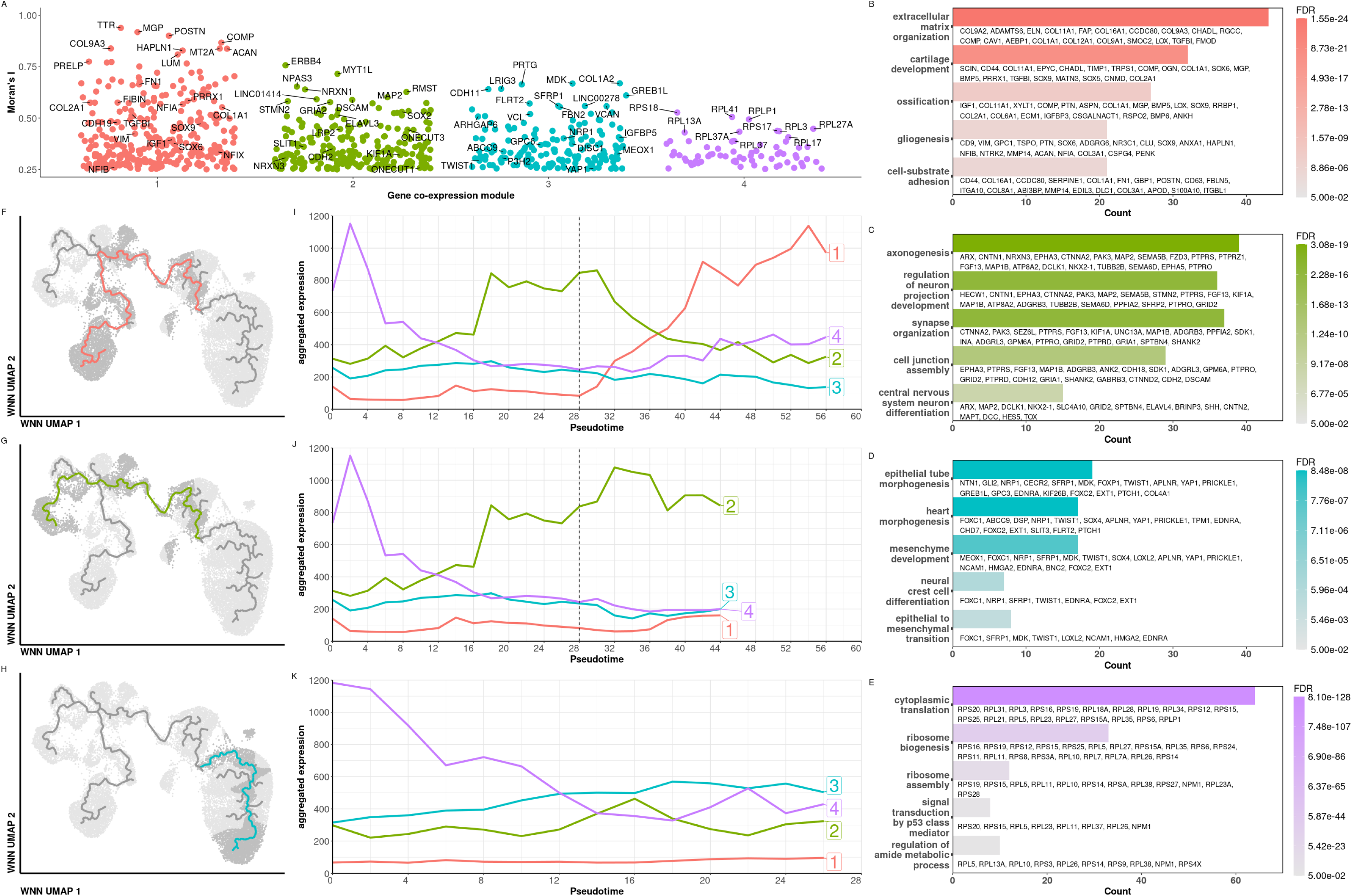
Pseudotime Analysis. **A**) Gene co-expression modules determined by clustering of genes with high autocorrelation (FDR < 0.001 & Moran’s I > 0.25) across all cells (louvain resolution = 0.00002). **B-E**) Bar plot of gene enrichment analysis for each module. Gene Ontology: Biological Process was used, and maximum 20 genes for every ontology included below the bar. Colours correspond to those in Figure 4A module-1 (B), module-2 (C), module-3 (D), module 4 (E). **F-H**) WNN UMAP projections of timepoints day-6, day-8, day-14, day-21, day-49 showing the trajectories highlighted with colour. Cells that contribute to these trajectories are indicated in a darker shade of grey. The trajectories are: Trajectory-1 (F, red), Trajectory-2 (G, green), and Trajectory-3 (H, blue). **I-K**) Aggregated expression of gene co-expression modules with colours and labels that correspond with Figure 4A over pseudotime with bins of 2 units for each trajectory. The dotted line indicates the day 21 bifurcation point. The trajectories are: Trajectory-1 (I), Trajectory-2 (J), andTrajectory-3 (K).

To next integrate the dynamics of these aggregated co-expression modules, we plotted them over pseudo-time along the three trajectories. **Fig. 3I** outlines the aggregated expression of the modules over pseudo-time of the chondrogenic **Trajectory-1**. It showed that expression of module-2 genes (green line) was prominent prior to the bifurcation point at day-21 (pseudo-time unit 28, vertical dotted line) whereas a switch between module-1 and module-2 occurred around pseudo-time unit 38, with prominent aggregated expression of module-1 until the chondrogenic terminus at pseudo-time unit 56. In **Fig. 3J**, it is shown that module-2 genes across the neurogenic **Trajectory-2** remained prominent until the terminus at pseudo-time unit 45 and was not overtaken by Module-1. Finally, as outlined in **Fig. 3K**, we observed that the mesenchymal neuro-crest module-3 genes had a relative high aggregated expression until the terminus of **Trajectory-3** at pseudo-time unit ∼28. A common feature in all three trajectories was the high aggregated expression of Module-4 genes at the earliest pseudo-timepoints that quickly drop off as pseudo-time progresses, corresponding to a high degree of translation and possible proliferation. Together these data demonstrate that chondrogenic cell-fate commitment emerges from the neural tube rather than the neural crest development.

### Characterizing transcriptome landscapes at day-21 divergent cell-fate trajectories

To characterize the molecular landscape of cells that constitute the divergent chondrogenic **Trajectory-1** (‘on-target’) and neurogenic **Trajectory-2** (‘off-target’) cell-fates, we first defined distinct clusters around the day-21 bifurcation by Louvain clustering (**Supplementary Fig. 3A**). In doing so, three cell clusters on the trajectory were defined (**Fig. 4A**), a pre-bifurcation state (cluster 10), post-bifurcation chondrogenic cell-fate state (‘on-target’ Trajectory-1, cluster 15), and post-bifurcation neurogenic cell-fate state (‘off-target’ Trajectory-2, cluster 8), together covering a relatively small pseudo-time window of 10 units. To identify distinct differences right after the day-21 bifurcation for Trajectory-1 and Trajectory-2, we performed differential expression analysis of the module genes. For the ‘on-target’ chondrogenic Trajectory-1, differential expression analysis of Module genes resulted in 180 (23%) significant differentially expressed genes (**Fig. 4B, Supplementary Table 6**) of which 134 (74%) were upregulated and 46 (26%) were downregulated. Among the upregulated genes we recognized 101 (75%) Module-1 genes (**Fig. 4B**, red dots) with notable highly significantly upregulated chondrogenic genes such as *FN1, MGP, DCN, FIBIN, ACAN,* and *IGFBP7*. On the other hand, differential expression of the ‘off-target’ neurogenic Trajectory-2, genes, showed 291 (38%) differentially expressed genes (**Fig. 4C, Supplementary Table 7**) with 175 (60%) upregulated and 116 (40%) downregulated genes. Among these upregulated genes we recognized 139 (79%) Module-2 genes (**Fig. 4C**, green dots) with notable, highly significant upregulated neurogenic genes such as *NRXN1, MYT1L,* and *STMN2*.

**Figure 4:**
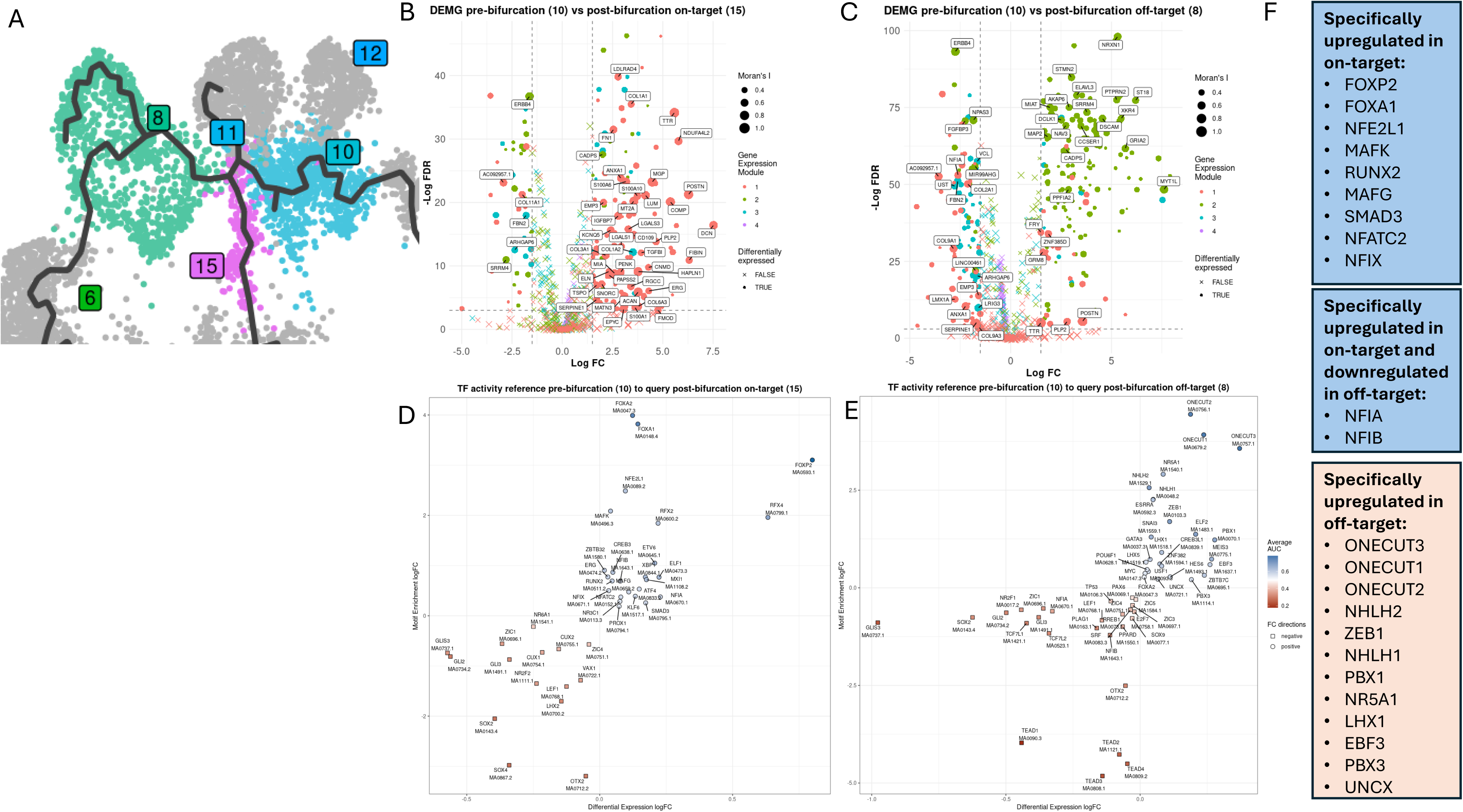
: Differential Analyses around trajectory bifurcation at day 21. **A)** WNN UMAP projection of days 6-49 of differentiation with inferred trajectory in black, clustered to fit around the bifurcation point at day-21. Clusters are labeled and cell colour corresponds to labels. Highlighted clusters represent pre-bifurcation (cluster-10), post-bifurcation ‘on-target’ (cluster-15) and post-bifurcation ‘off-target’ (cluster-8). **B)** Volcano plots showing differential expression of module genes between pre-bifurcation (cluster-10) as reference and post-bifurcation ‘on-target’ (cluster-15) as query and **C)** between pre-bifurcation (cluster-10) as reference and post-bifurcation ‘off-target’ (cluster-8) as query. Differentially expressed module genes have FDR < 0.05 and a log FC > |1.5| and coloured by module. Genes with Moran’s I score ≥ 0.5 are labeled by gene name. **D)** Scatter plot showing differential transcription factor activity assay (TFAA) between pre-bifurcation (cluster-10) as reference and post-bifurcation ‘on-target’ cluster-15 as query and **E)** between pre-bifurcation cluster-10 as reference and post-bifurcation ‘off-target’ cluster-8 as query. Each point is labeled with the TF gene name and TF binding motif name underneath. The shape indicates the direction of activity change congruent between differential expression and differential motif enrichment. **F)** Exclusive ‘on target’ and ‘off target’ transcription factors. See also Supplementary Figure 3B.

To identify transcription factors that direct these chondrogenic or neurogenic cell-fate commitment, we performed transcription factor activity assay (TFAA) in a differential manner in both the ‘on-target’ and ‘off-target’ direction. Upon plotting differential expression and differential enrichment of TF binding motifs for the chondrogenic ‘on-target’ trajectory (**Fig. 4D**), we showed that the activity of TFs *FOXA1*, *FOXA2*, *FOXP2*, and *RFX4* had the largest increase, and *GLIS3*, *GLI2*, *SOX2*, *SOX4*, and *OTX2* had the largest decrease. On the other hand, TFAA for in the neurogenic ‘off-target’ trajectory (**Fig. 4E**) we showed that the activity of TFs *ONECUT1*, *ONECUT2*, and *ONECUT3* had the largest increase, and *TEAD1, TEAD2, TEAD3, TEAD4,* and *OTX2* had the largest decrease in activity.

To prioritize potential TFs that are exclusively directing the chondrogenic ‘on-target’ or neurogenic ‘off-target’ cell-fate, we integrated a third differential TFAA, now comparing the neurogenic Trajectory-2 (cluster 8) with the chondrogenic Trajectory-1 (cluster 15). As outlined in **Supplementary Figure-3B**, we identified TFs such as *FOXP1, NFIA, SMAD3* and *NR2F1* with the largest upregulation in activity in the chondrogenic direction and TFs such as *ONECUT2, ZEB1, ONECUT1, PBX1, PBX3, and ONECUT3,* with the largest upregulation in the neurogenic direction. By subsequently overlapping the results of the three TFAAs (**Supplementary Figure-3C**) we identified exclusive TFs. As shown in **Supplementary Figure-3C** and **Fig. 4F**, *FOXP2*, *FOXA1*, *NFE2L1*, *MAFK*, *RUNX2*, *MAFG*, *SMAD3*, *NFATC2*, *NFIX*, *NFIA*, and *NFIB* became exclusively and significantly more active in the chondrogenic post-bifurcation trajectory (Trajectory-1). Conversely, *ONECUT3*, *ONECUT1*, *ONECUT2*, *NHLH2*, *ZEB1*, *NHLH1*, *PBX1*, *NR5A1*, *LHX1*, *EBF3*, *PBX3*, and *UNCX* became exclusively and significantly more active in the neurogenic post-bifurcation trajectory (Trajectory-2). Noteworthy is that among the chondrogenesis specific TFs, *NFIA* and *NFIB* showed the strongest oppositional regulation between cell-fates hence may play a pronounced role in chondrogenic cell-fate commitment.

### Characterizing temporal transcription factor activity over pseudotime

To obtain insight into the temporal dimension of the exclusively and significantly more active transcription at the day-21 bifurcation (**Fig. 4F**), we next plotted their expression patterns over pseudo-time. In the chondrogenic ‘on-target’ direction (**Fig. 5A**) it was shown that expression of *NFIA* and *NFIB* follow a very similar patterns along pseudo-time of Trajectory-1, though *NFIA* had considerably higher levels of expression. Notable is also that the expression peak for both *NFIA* and *NFIB* was located past the bifurcation point at day-21 (i.e. 28 units of pseudo-time), indicating their effects observed in the regulatory networks increased further. Moreover, *NFIA* and *NFIB* were especially prevalent in day-49 pre-terminal chondrogenic clusters while *NFIX* was most expressed in the terminal chondrocyte like cluster (**Fig. 5E**). This indicated that these Nuclear Factor I genes play a driving role in chondrogenic cell-fate commitment. In the neurogenic ‘off-target’ direction (**Fig. 5D**) it was shown that expression of *ONECUT1, ONECUT2,* and *ONECUT3* also followed similar patterns along pseudo-time of trajectory-2. Notable is that the expression of *ONECUT3* showed a clear peak past the bifurcation point at day-21. Moreover, *ONECUT1, ONECUT2*, and *ONECUT3* were prevalent in the off-target neurogenic Trajectory-2 specifically at day 21 (**Fig. 5E**).

**Figure 5:**
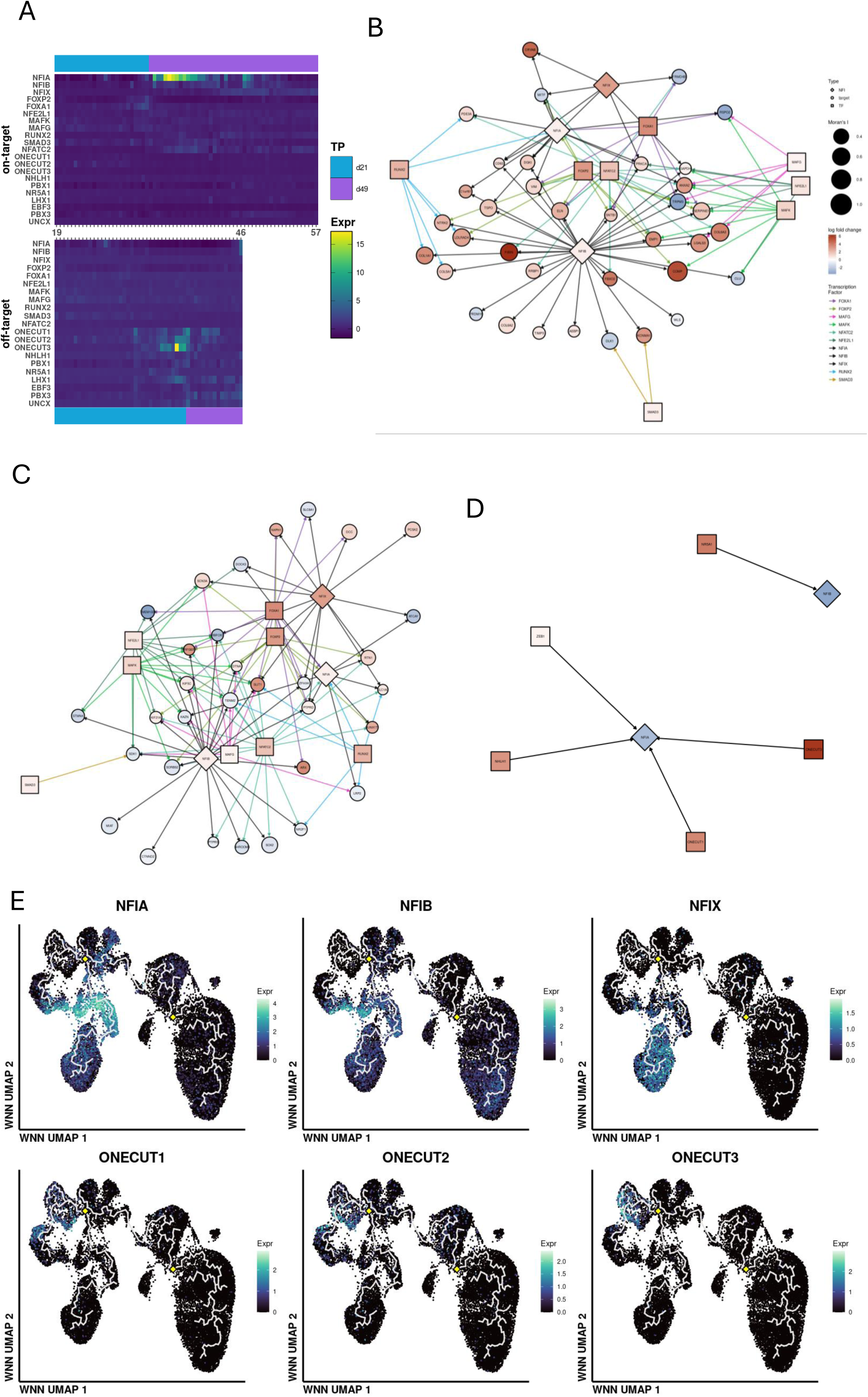
Transcription factor dynamics. **A-D**) Nodes are shaped by category, diamond for Nuclear Factor-1 genes, round for target module genes, and square for other chondrogenic exclusive transcription factors (TF). Nodes are coloured by differential expression between pre-bifurcation cluster 10 (reference) and post-bifurcation ‘on-target’ cluster 15 (query). Node size is set by Moran’s I score. Edges show the target interaction between transcription factors and target genes by the direction of the arrow, edges are coloured by the node of origin. **A**) Transcription factor-target network showing the effect on all module genes targeted by Nuclear Factor-1 genes in the chondrogenic ‘on-target’ direction, Adapted from **Supplementary Figure 5**, as are 6B and 6C. **B**) Transcription factor-target network showing the effect on all module-1 genes targeted by Nuclear Factor-1 genes and co-factors in the chondrogenic ‘on-target’ direction. **C**) Transcription factor-target network showing the effect on all module-2 genes targeted by Nuclear Factor-1 genes and co-factors in the chondrogenic ‘on-target’ direction. **D**) Transcription factor-target network of the neurogenic ‘off-target’ direction. Here showing the transcription factors which target Nuclear Factor-1 genes NFIA and NFIB. **E**) Heatmap of SCT normalized expression over pseudotime in bins of 0.5 units in the chondrogenic ‘on-target’ Trajectory-1 and the neurogenic ‘off-target’ Trajectory-2 for six transcription factors of interest. **F**) WNN UMAP projection of days 6-49 of differentiation with trajectories indicated in light-grey, coloured by the SCT normalized expression of the gene of interest. From left to right, top to bottom; *NFIA*, *NFIB*, *NFIX*, *ONECUT1*, *ONECUT2*, *ONECUT3*.

### Mapping transcription factor – target networks by cis-co-accessibility

To map transcription factor gene-target networks that drive chondrogenic cell fate commitments at the day-21 bifurcation point, we implemented cis-co-accessibility of differential enrichment of TF binding motifs (TFAA, Fig. 4E-F) and differentially expressed module genes (Fig. 4C-D). In the chondrogenic, ‘on-target’ Trajectory-1, we identified cis-co-accessibility for *NFIA*, *NFIB*, and *NFIX* with 37 differentially expressed chondrogenic Module-1 genes (**Fig. 5A**) and 32 differentially expressed neurogenic Module-2 genes (**Fig. 5B**). Notable herein is that 26 out of 37 Module-1 genes were upregulated whereas 18 out of 32 Module-2 genes were downregulated, demonstrating the directive role of *NFIA*, *NFIB*, and *NFIX* in branching the chondrogenic Trajectory-1 from the neurogenic Trajectory-2. More specifically, among the upregulated chondrogenic Module-1 genes, we showed cis-co-accessibility between *NFIB* with *COMP, FIBIN,* and *COL1A1,* and between *NFIA* and *NFIB* and *VIM*. On the other hand, among the down regulated Module-2 genes we showed cis-co-accessibility between *NFIB* and *NFIA* and the neuroepithelial markers *SOX2* and *LRP2*, respectively. Notable was also that cis-co-accessibility existed between *NFIA* and other TF driving chondrogenic cell fate, namely *FOXA1*, *FOXP2*, *NFATC2*, and *RUNX2*. This created triangular connections (**Fig. 5A**) that allowed fortification of transcription factor - target regulation. For the neurogenic ‘off-target’ Trajectory-2, cis-co-accessibility network showed that *NFIA* is actively downregulated by *ONECUT1*, *ONECUT3, ZEB1,* and *NHLH1*, (**Fig. 5C**, and **Supplementary Table 10**) whereas, *NFIB* was downregulated by *NR5A1* (**Fig. 5C**). Together, our data demonstrated highly interactive regulatory TF target networks driving cell fate commitment at the day-21 bifurcation with *NFIA* and *NFIB* and *ONECUT1 ONECUT3* as important players. Herein, *NFIA* and *NFIB* activity appeared mandatory for branching the chondrogenic (Trajectory-1) from the neurogenic (Trajectory-2) cell fate commitment.

### Nuclear Factor 1 binding motifs direct chondrogenic cell-fate commitment

To investigate the *NFIA* interactive regulation more closely, we next focused on the cis-co-accessibility networks (**Fig. 6A**, track Links) at genomic region of the *NFIA* promoter. Specifically, we focused on the promotor of the most reliable *NFIA* isoform (ENST00000403491.8, chr1:61082561-61462788, **Fig. 6A**, track Genes). Subsequently we mapped transcription factor binding motifs from both the chondrogenic drivers *FOXA1*, *FOXP2*, *NFATC2*, and *RUNX2,* but also the neurogenic drivers *ONECUT1*, *ONECUT3, ZEB1* and *NHLH1* (**Fig. 6A**, track ‘TF binding). Finally we plotted normalized signals of chromatin accessibility of the pre-bifurcation (cluster 10), post-bifurcation chondrogenic Trajectory-1 (cluster 15), and post-bifurcation neurogenic Trajectory-2 (cluster 8) and their corresponded *NFIA* expression (**Fig. 6B**). Notable in **Fig. 6A** were the high normalized chromatin signals of Peak-2 (chr1:61049780-61050726), Peak-3 (chr1:61076610-61077482), and Peak-4 (chr1:61077731-61078553), that were significantly differentially accessible between pre-bifurcation cluster 10 and neurogenic ‘off-target’ cluster-8 (**Fig.6A** grey highlight). As shown in the TF-binding track, these differential chromatin peaks, were host to transcription factor binding sites of the exclusively upregulated chondrogenic ‘on-target’ transcription factors *FOXP2* (Peak-3) and *NFATC2* (Peak-3 and Peak-4), indicating active regulation of *NFIA* in Trajectory-1 at the day 21 bifurcation. The complex orchestration of this region is further illustrated by the observation that also the neurogenic ‘off-target’ transcription factors *NHLH1*, and *ZEB1* (Peak-2 and Peak-3) appear to have binding sites here. However, since these are not upregulated in the chondrogenic ‘on-target’ Trajectory-1, they will not interfere with *NFIA* expression regulation.

**Figure 6:**
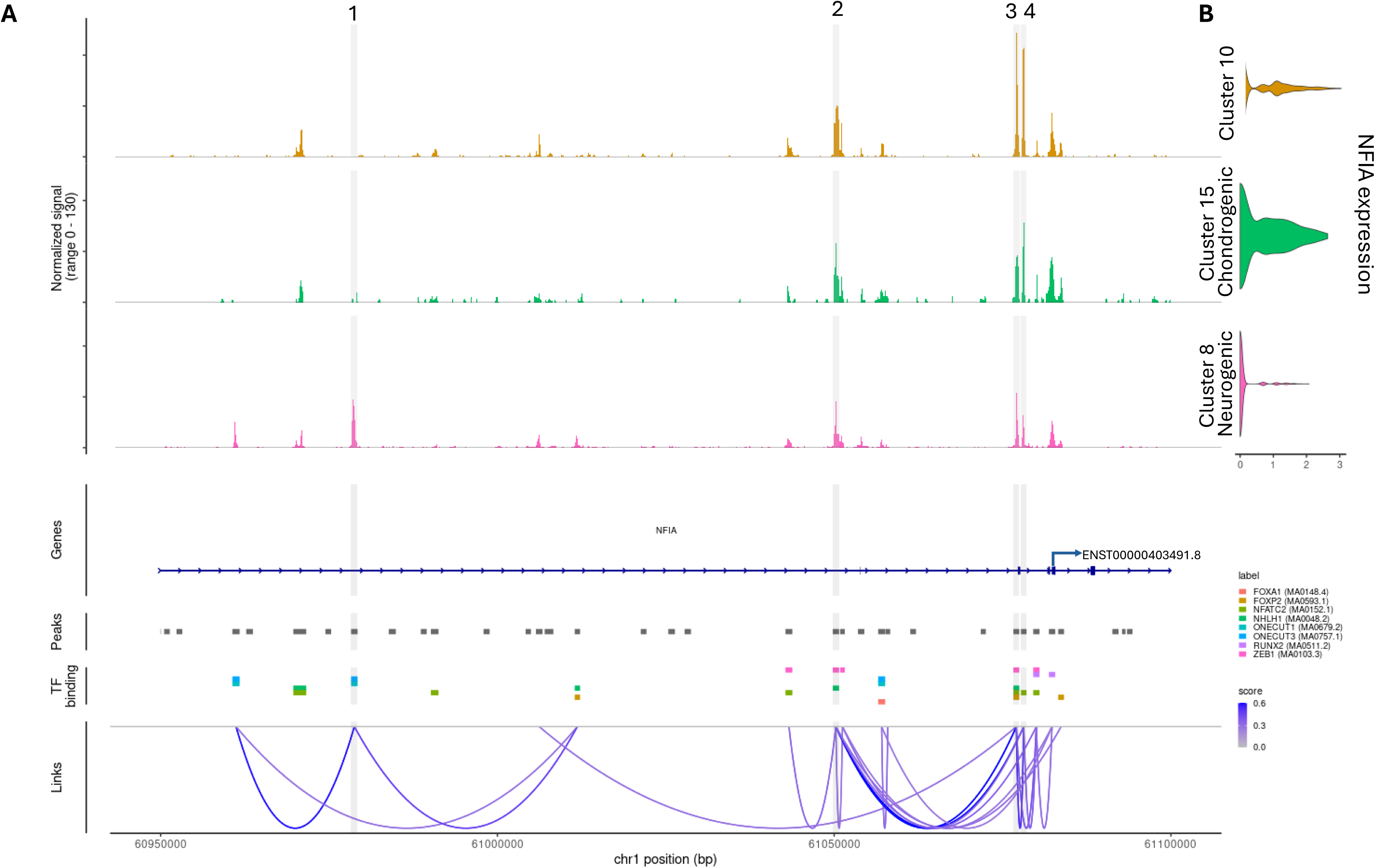
Coverage plot of the NFIA promotor region chr1:60950000-61100000 (hg38), adapted from **Supplementary Figure 5**. **A**) The first three tracks show normalized signal (fragment count) across the range of interest for the pre-bifurcation cluster 10 (orange), chondrogenic post-bifurcation ‘on-target’ cluster 15 (green), neurogenic post-bifurcation ‘off-target’ cluster 8 (pink). The fourth track shows the gene body, here indicated is the transcription start site of NFIA isoform ENST00000403491.8 (chr1:61082561-61462788). The fifth track shows peaks called from fragments. The sixth track shows transcription factor binding sites in peaks that make up CCAN nr. 520 in the region of interest. Only TFs that target NFIA are included for the chondrogenic trajectory (*FOXA1*, *FOXP2*, *NFATC2*, *RUNX2*) and the neurogenic trajectory (*NHLH1*, *ONECUT1*, *ONECUT3*, *ZEB1*). The seventh track shows the links between peaks that makes up cis-co-accessibility network nr. 520 in the region of interest colored by co-accessibility score. **B**) Violin plots showing SCT normalized expression of *NFIA* in the clusters of interest corresponding to the first 3 tracks of 7A: pre-bifurcation cluster 10 (orange), chondrogenic post-bifurcation ‘on-target’ cluster 15 (green), neurogenic post-bifurcation ‘off-target’ cluster 8 (pink).

Finally, an inverse situation unfolded at identified network peaks located more distally from the *NFIA* promotor region Peak-1 (chr1:60978278-60979241) where accessibility was significantly increased in the neurogenic ‘off-target’ cluster-8. This network peak only contained binding sites for exclusively upregulated neurogenic ‘off-target’ transcription factors *ONECUT1* and *ONECUT3*, which therefore could be part of the observed downregulated *NFIA* transcription in the neurogenic ‘off-target’ direction by these TFs. Taken together, this data shows that *FOXP2*, *RUNX2*, but especially *NFATC2* bind the promotor region to upregulate *NFIA* expression and that *NHLH1* and *ZEB1* bind the promotor region to downregulate *NFIA* expression with *ONECUT1* and *ONECUT3*.

### Exogenic NFIA promotes chondrogenic differentiation at day 21

Next we experimentally validated the driving role of NFIA in chondrogenesis, by transiently upregulating *NFIA* expression around the day-21 point of bifurcation (**Fig. 7A**). Herein we focussed on identified direct targets of *NFIA* (Fig. 5B) and confirmed, as shown in **Fig. 7B**, that *NFIA* directly upregulated the selected Module-1 target genes *VIM* and *CRYAB*. Unexpectedly however, transient *NFIA* upregulation also upregulated the Module-2 gene *ATCAY*. Moreover, as shown in **Fig. 7C**, the strong and significant upregulation of the prominent chondrogenic Module-1 genes *MGP* and *COMP*, further confirms the chondrogenic driving force of *NFIA.* Even more, we showed that the transient upregulation of *NFIA* around the day-21 point of bifurcation was sufficient to enhance neo-cartilage organoid formation both in size and matrix deposition as reflected by the glycosaminoglycan-rich cartilaginous tissue (Alcian blue staining, **Fig. 7C**) but also by the opacity of intact organoids under bright field light microscope (**Supplementary Figure 6**). Together these data confirms that transient upregulation of *NFIA* directs chondrogenic cell-fate commitment towards high quality neocartilage organoids.

**Figure 7:**
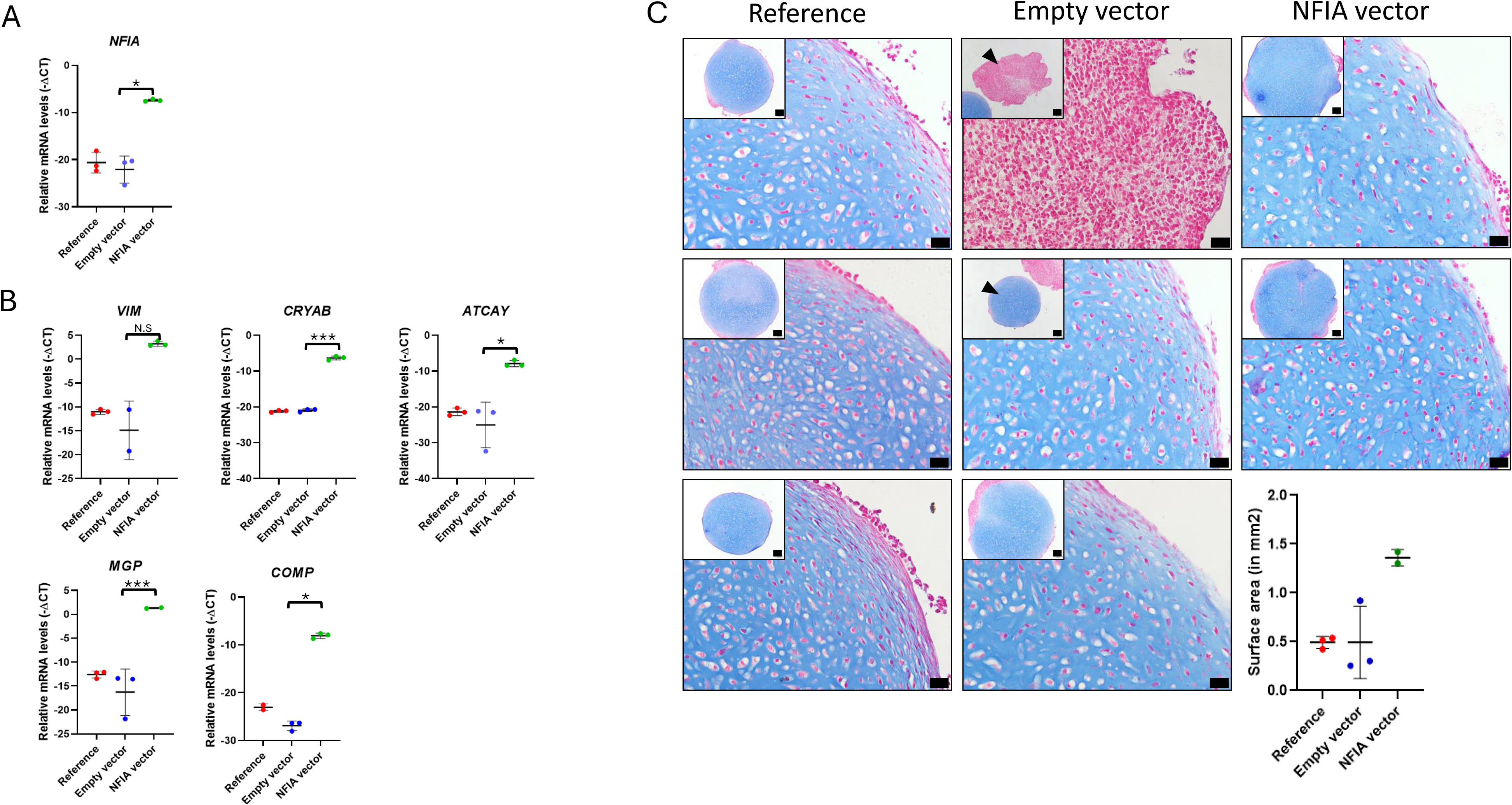
NFIA transfection experiment. **A)** From left to right, top to bottom: gene expression minus delta CT is given for *NFIA*, *VIM*, *CRYAB*, *ATCAY*, *NFIB*, *COMP*, *FIBIN*, and *MGP*. **B)** Brightfield Microscopy images of day 35 neocartilage organoid coupes with alcian blue staining. From left to right are conditions: Reference in triplo, Empty vector in triplo (arrows point to whole and separate organoids), and NFIA vector in duplo. Per panel: large is 40x magnification (20 µm black bar), small top left is 10x magnification (100 µm black bar). The bottom right panel shows a surface area quantification of each coupe.

## Discussion

To map hiPSC chondrogenic differentiation and understand the regulatory mechanism behind chondrogenic cell-fate decisions, we performed multi-modal (mRNA and ATAC) simultaneous single-nucleus sequencing at 7 timepoints during an established hiPSC differentiation to living non-hypertrophic neocartilage organoids. By leveraging temporal and multimodality aspects of our single-nucleus dataset, we demonstrated that the chondrogenic trajectory initiated at day-6 from a neurogenic development that subsequently branched off at day-21 to an ‘on-target’ chondrocyte (Trajectory-1) cell fate. The solid ‘on-target’ chondrogenic branch that emerged at day-21 was marked by an abundant expression of chondrogenic markers such as *COL2A1, ACAN, MGP,* and *COMP*. Transcription factor activity analyses, demonstrated that activity of *NFIA*, *NFIB*, and *NFIX* of the nuclear factor-1 family, critically drove the chondrogenic split by directly targeting prominent chondrogenic genes such as *COMP, FIBIN,* and *VIM*, but also downregulating neuroepithelial markers such as *SOX2*, and *LRP2*. In turn, exploring the cis-co-accessibility networks in the genomic promotor region of *NFIA*, subsequently showed highly interactive regulatory TF target networks involving for example *FOXP2, NFATC2,* modulating the chondrogenic action of *NFIA* further. The direct targets of *NFIA* were experimentally validated, resulting in high quality neocartilage organoids that abundantly expressed chondrogenic genes. Together, our data showed that in the hiPSC chondrogenic differentiation, *NFIA* and *NFIB* appear to be mandatory for branching the chondrogenic from the neurogenic cell fate commitment towards non-hypertrophic high quality neocartilage organoids. As such offering a rational basis to advance engineering of high-fidelity living cartilage constructs for regenerative medicine.

Owing to the fact that both mRNA expression and open chromatin modalities were extracted from a single nucleus at 7 timepoints during chondrogenic differentiation, we could reliably map dynamic changes in transcription factor activity across critical (pseudo)-timepoints. Taking this further, we could seamlessly combine ATAC-based CCAN constructions with expression data as the source remained the same. This allowed us to, instead of identify a core set of transcription factors, generate a gene regulatory network and elucidate TFs that actually drive cell fate decisions at the point of bifurcation at day 21. It is through these mechanisms that we also demonstrated that the ‘off-target’ neurogenic branch continued down with the neurogenic cell-fate commitment, expressing markers such as *MYT1L* and *NRXN1*, whereas, TFAA analyses showed that *ONECUT1* to *ONECUT3* were most prominent in maintaining neurogenic cell-fate.

Previously, it was indicated that, despite neurogenic ‘off-target’ cells, the applied chondrogenic hiPSC differentiation protocol followed the well-known canonical development of cartilage via the mesodermal route.^8^ A route that is, however associated with aberrant hypertrophic, terminal maturing chondrocytes.^4,10^ Since we only changed handling of chondroprogenitors at day-14 compared to Wu et al.^8^, picking aggregates instead of dissociating and re-pelleting the chondroprogenitor colonies, we had assumed previously that we also followed mesodermal lineage differentiation to human induced chondrocytes (hiCHOs).^5^ It was revealed however, in the current study that the ‘on-target’ chondrogenic trajectory from day-6 to day-21 retain an expression profile in line with neuroepithelial cells of the neural tube and thereby the central nervous system development (Module-2 genes). Notable is also that Trajectory-3, which started at day-6 and terminated at day-8, appeared to fulfil the neural crest cell to mesenchyme trajectory via epithelial to mesenchymal transition, as marked by Module-3 genes such as *FLRT2*, *LRIG3*, and *DISC1*, that was not present in day-14 cells. We advocate, that the picking of chondroprogenitor colonies at day-14 is essential in committing to a high quality, non-hypertrophic chondrocyte cell-fate via neural tube lineage. The fact that our hiCHOs had very low expression of the hypertrophic markers, such as *COL10A1*, *TNFRSF11B*, or *SPP1*, is in line with findings that as ectoderm derived cranial neocartilage is resistant to endochondral ossification, unlike mesodermal neocartilage.^11–14^ Taken together, here have characterised a novel differentiation route that forgoes mesodermal and neural crest cell fates, emerging from the neuroepithelial cells of the neural tube.

At day-21, one week after transferring the aggregates to 3D suspension cultures a in regular chondrogenic differentiation medium containing TGFB1, a solid bifurcation was observed that resulted in a chondrogenic (Trajectory-1) and neurogenic (Trajectory-2) trajectory both ending at day-49. As such a mixed population of chondrocytes and neurogenic cells occur at the day-49 of hiPSC-chondrogenic differentiation that could explain the previously observed results of relative low expression of the chondrogenic markers *COMP*, *MGP*, and *VIM* and detectable expression of the neuronal marker *KIF1A*.^5,8^ An item that is open for discussion is whether Trajectory-2 cells represent a stable ‘off-target’ developmental trajectory to neurogenic cells, a dynamic maturing system with remaining chondrogenic developmental potential, or a supporting cell population for chondrogenesis. The fact that at the terminus of Trajectory-2, *NFIA* expression becomes detectable suggests greater plasticity towards a chondrogenic cell-fate than expected of a matured cell population.

There is literary precedent to suggest Nuclear Factor-I genes can drive chondrogenic cell fate commitment in addition to neuronal development. Firstly, Nuclear Factor-I genes have long been associated with development of the central nervous system and in particular the cerebellum, where they play a role in neuro-progenitor and glial cell-fate commitment^15–17^ whereas *Nfia* and *Nfib* knock-out mice showed a delayed central nervous system development phenotype.^18^ Second, recent studies have shown that *NFIA* and *NFIX* play a role in the development of cartilage and *NFIA* in particular is involved in cartilage maintenance through fatty acid metabolism.^19–21^ Finally, dysfunctional *NFIX* is known to cause, Marshall-Smith syndrome and Malan syndrome, that both manifest with neurodevelopmental delay but also with a short stature and accelerated bone maturation. The latter, further supporting a role for Nuclear Factor-I genes in maintaining a chondrocyte phenotype in humans.^22–24^

Collagen type 2 is a prominent constituent of cartilage and as such has previously served as important marker for chondrogenic differentiation.^25^ Here we show that *COL2A1* expression, however, fluctuates during chondrogenic differentiation, hence may not be the best marker. Particularly, *COL2A1* expression was low early along the ‘on-target’ Trajectory-1 (starting at day-6 and progressing through day-14) while being high in cells committed along Trajectory-3, i.e. cells that do not progress to hiCHOs. On the other hand Vimentin, showed a steadily increasing in expression along Trajectory-1 right up to the terminal cluster at day 49. We propose, therefore, Vimentin as potential novel marker for progression of hiPSC chondrogenic differentiation. On a different note, there are no cells at day-49 which are similar to hiPSCs indicating a very low likelihood to the formation of any teratomas upon using this protocol for regenerative cartilage tissue transplantation.

A perceived weakness of our study is the fact that, in our focus to explore chondrogenic cell fate, we generated data in the early stages of differentiation at day-0, day-3, and day-6, only. Therefore, in this early phase of differentiation, cell populations were disconnected from each other and the proceeding day-6 continuum and early trajectories could not be inferenced. Moreover, this implies that we could not identify day-6 pre-bifurcation cell states. Additionally, albeit of good quality, the number of cells sequenced at day-14 were relatively low which hampered robust mapping of the contribution of day-14 cells to the respective clusters. Finally, we used hiPSC chondrogenic differentiation as analogous model to map cartilage development. Notably in this respect is that Jachim et al.^27^ recently showed similar transcription factors playing a role in in human foetal cartilage development.

Together, we generated here, for the first time, a temporal and multimodal single-cell dataset across hiPSC-chondrogenic differentiations. The temporal and multimodal aspects of this dataset allowed us to reliably generate dynamic gene regulatory networks and unravel how they drive cell fate decisions at the point of bifurcation at day 21. Herein, *NFIA* and *NFIB* were found to critical modifiable transcriptional regulators that can now be leveraged to implement regenerative cartilage therapies from hiPSCs.

## Materials and Methods

### hiPSC culture

The hiPSC line LUMC0004iCTRL10 is male skin fibroblast derived without any known diseases and has been registered with the Human Pluripotent Stem Cell Registry^28^. The hiPSC line was generated with approval from the Leiden University Medical Center Ethical Committee under P13.080. Human iPSCs were cultured on Vitronectin-XF (STEMCELL Technologies) at 37⁰C, 5% CO_2_ and were refreshed every 24 hours with TeSR-E8 medium (STEMCELL Technologies).

### hiPSC differentiation

Chondrogenic differentiation of hiPSCs was completed as described before^10–13^. The hiPSCs were passaged and expanded for 3 days in 2D on Vitronectin-XF before initiation of the differentiation in basal differentiation medium (50% IMDM GlutaMAX (Gibco-ThermoFisher Scientific) and 50% Ham’s F-12 Nutrient Mix (Sigma-Aldrich), supplemented with 1% Chemically defined lipid concentrate (Gibco-ThermoFisher Scientific), 1% ITS+ Premix Universal Culture Supplement (Corning), 0.2% Penicillin/Streptomycin (Gibco-ThermoFisher Scientific), and 450µM 1-Thioglycerol (Sigma-Aldrich)). Cells were refreshed every 24 hr with different morphogens. On Day 0, basal medium was supplemented with 30ng/ml Activin A (Miltenyibiotec), 4μM of CHIR99021 (Tocris), and 20ng/ml recombinant human fibroblast growth factor (basic, Peprotech). On day 1, cells were refreshed with basal medium supplemented with 2μM SB-505124 (Tocris), 3μM CHIR99021, 20 ng/ml recombinant human fibroblast growth factor, 4μM Dorsomorphin dihydrochloride (Tocris). On day 2 basal medium was supplemented with 2μM SB-505124, 1μM Wnt-C59 (Tocris), 500nM PD173074 (Tocris), and 4μM Dorsomorphin. On days 3-5, basal medium was supplemented with 2μM Purmorphamine (Tocris) and 1μM Wnt-C59. And during days 6 – 13 basal medium was supplemented with 20 ng/ml human bone morphogenetic protein 4 (premium grade, Miltenyibiotec). Prior to refreshing the medium, cells were washed with plain IMDM GlutaMAX and Ham’s F-12 Nutrient Mix (ratio 1:1), and 0.2% Penicillin/Streptomycin.

The differentiated cells were then transferred from 2D cell culture to suspension cell culture in a 15ml falcon tube for the second stage of chondrogenic differentiation. Cells were selected based on the formation of dense 3D aggregate structures in the 2D dish. Aggregates were scraped-off gently from the plate and transferred into suspension culture where they were then centrifuged at 300g for 5 minutes. Media was then aspirated from the cell pellet and 0.5ml chondrogenic differentiation medium (CDM) consisting of DMEM/F-12 GlutaMAX (Gibco-ThermoFisher), supplemented with 1% ITS+, 55μM 2-mercaptoethanol (Gibco-ThermoFisher), 100nM dexamethasone (Sigma-Aldrich), 1% MEM Non-Essential Amino Acids (Gibco-ThermoFisher), 0.2% P/S, 50μg/ml (+)-Sodium L-ascorbate (Sigma-Aldrich), 40μg/ml L-proline (Sigma-Aldrich), and 10ng/ml human recombinant TGF-β1 (HEK293 derived, Peprotech). Aggregates were cultured at 37⁰C, 5% CO2 and CDM was refreshed every 3-4 days until collection of the neocartilage for nuclei isolation.

### hiCPC dissociation and nuclei isolation

Neocartilage were dissociated using a two-phase extracellular matrix (ECM) digestion protocol. Phase one involved suspending the organoids (3 per reaction) in 1.5ml chondrogenic differentiation medium (CDM) containing 2mg/ml Pronase (Roche) and incubating at 37⁰C for 45 minutes. In phase two, organoids were centrifuged at 150g for 3 minutes and Pronase supplemented media was aspirated and replaced with 1.5ml CDM containing 1.5mg/ml Collagenase I (Antonides) for one hour incubation at 37⁰C followed by centrifugation at 150g for 3 minutes. Medium was aspirated, and dissociated cells were washed with PBS and filtered through a 70µm pore strainer to remove aggregates. Subsequently, cells were counted and tested for viability, and 10000 cells were then centrifuged at 150g for 3 minutes and resuspended in 1ml PBS.

Nuclei were isolated from the dissociated cells according to 10X Genomics recommendation^29^. In short, after centrifuging cells at 300g for 5 minutes at 4⁰C, PBS was removed and 100µl chilled lysis buffer (nuclease free water containing 0.1% NP40 substitute (Sigma-Aldrich), 0.01% digitonin (Invitrogen), 0.01M Tris-HCl (Invitrogen), 0.01M NaCl (JT Baker), 3mM MgCl2 (Boom), 1% bovine serum albumin (Sigma-Aldrich), 0.1% Tween-20 (Sigma-Aldrich), and 0.5U/ml RNAse inhibitor (Roche)) was added to the cells and incubated on ice for 5 minutes. Immediately after the incubation period 1ml washing buffer (nuclease free water containing 0.01M Tris-HCl, 0.01M NaCl, 3mM MgCl2, 1% bovine serum albumin, 0.1% Tween-20, and 0.5U/ml RNAse) was added to the nuclei in lysate by gentle mixing followed by centrifugation at 500g for 5 minutes at 4⁰C. Finally, the supernatant was removed, and the remaining nuclei resuspended in PBS containing 2% BSA in PBS for multi-modal sequencing.

### Multimodal single nucleus sequencing

Isolated nuclei were used to generate a single nucleus RNA and ATAC sequencing libraries using the 10X Genomics Chromium Next GEM Single Cell Multiome ATAC + Gene Expression Kit (PN-1000285) and Chromium Next GEM Chip J (PN-100023) according to the manufacturer’s protocol. snATAC-seq libraries were sequenced on an Illumina NextSeq2000 platform and snRNA-seq libraries were sequenced on a NovaSeq6000 platform with v1.5 chemistry. Samples were demultiplexed using bcl2fastq v2.20 and count tables were generated using the 10x Genomics Cell Ranger ARC software v2.0.1 and reference file refdata-cellranger-arc-GRCh38-2020-A-2.0.0.

### Quality control and pre-processing snRNA-seq and snATAC-seq data

Multimodal data was processed using Seurat v5.1.0^30^ and Signac v1.13.0^31^. Quality was determined by correcting for low feature count, doublets and percentage of mitochondrial or ribosomal RNA features in RNA and low feature count, doublets, and percentage peak fragments in ATAC (**supplementary figure 1**). First, in the RNA modality, empty cells were found by low feature count and eliminated under the 15.8-20^th^ percentile, exception for day-3 with 1^st^ percent as attributed to the high cell count. Doublets were eliminated by removing cells in the 99^th^ percentile of features. High mitochondrial and Ribosomal percentage with low feature count indicates genomic RNA damage, 95-99^th^ percentile by mitochondrial percentage were removed and the 85-90^th^ percentile by ribosomal percentage were removed, exception for day 21 where ribosomal cutoff was at 77^th^ percentile. Cell Cycle status for each cell was determined by S-score and G2M-score as provided by Seurat. Normalisation was done using SCTransform^32^, regressing out S-score, G2M-score, percentage mitochondrial RNA, and percentage ribosomal RNA. Initial dimensionality reduction was done using PCA on all genes. Second, in the ATAC modality, empty cells were found by low peak region fragments cutting off below the 22.8^th^ percentile. Doublets were eliminated by too high peak region fragments, cutting off at the 98.17^th^ percentile. The majority of reads should be in peaks (> 50%), we found that for each sample less than 6% of cells had less than 50% reads in peaks, with exception of day-14 where this was 10%. Transcriptions Start Site Enrichment was remarkably consistent even at the lowest percentiles (< 1%) and nucleosome score was remarkably consistent even at high scores (> 99%), no cells were eliminated on the basis of these metrics. Lastly we used Signac recommendation for eliminating cells by blacklisted peaks ratio set at 5%. Only cells that were sufficiently high quality in both modalities were included.

### Multimodal clustering analysis

Modalities were combined in Seurat using weighted nearest-neighbour network (WNN) analysis^3^. Then, uniform manifold approximation and projection (UMAP) was done for the combined modalities and clustered using K-nearest neighbour (KNN) and Louvain algorithm (LSM). Clustering resolution was selected based on highest mean silhouette width, lowest cluster size of at least 50 cells and visual confirmation.

### Trajectory Inference

Trajectory inference was performed using monocle 3 v1.3.7^33^in accordance with the available vignette. For this the UMAP projection from Seurat was used instead of recreating one.

### Gene autocorrelation and co-expression module construction

Gene co-expression modules were calculated using monocle 3 following the steps as outlined in the vignette. In short, genes get a moran’s I score based on the autocorrelation along the trajectories.

Thereby giving an expression specificity score per gene. We then used the most specific genes Moran’s I > 0.25 to cluster genes by expression across all cells day-6 to day-49 by Louvain clustering.

### Pathway analysis

Gene Ontology: Biological Process (GO:BP) overrepresentation analysis (ORA) was performed using clusterProfiler v4.10.0^29^ using all expressed genes in the dataset as a background or universe.

#### Pseudo-time analysis

Pseudo-time analysis was done using monocle 3, following the steps as outlined in the vignette. In short, cells are ascribed to nodes along a trajectory were isolated by trajectory, ordered by pseudo-time and split into bins of 0.5 in pseudo-time, aggregated expression for all genes in a module for each module was calculated and plotted over pseudo-time. Every bin of pseudo-time was ensured to have over 50 cells. Aggregated expression was not scaled as in the vignette.

### Differential Expression Analysis

Differential Expression analysis was done using DElegate to employ EdgeR v4.0.7^34^ for pseudoreplicate pseudobulk analysis on single cell data. Pseudoreplication was randomised as handled by Delegate. Differentially expressed genes were then selected based inclusion in co-expression modules to be differentially expressed module genes.

### Transcription Factor Activity Assay

Transcription Factor Activity Assay was done using Signac following the Signac vignette. For this the packages chromVAR v1.24.0^35^, JASPAR v0.99.10^36^, and presto v1.0.0 were used. The log fold change in differential gene expression and differential motif enrichment was used to determine a positive or negative change in activity by consensus. Finally, trajectory specific genes were determined by overlap in negative and positive change in activity depending on condition (**Supplementary Figure 3**).

### Transcription Factor-Target Networks

Transcription factor target networks were constructed from differentially active transcription factors specific for a trajectory and differentially expressed module genes as potential target. Target status was determined using cis-co-accessibility networks (CCAN) using cicero v1.3.9^37^. These were calculated over clusters 8, 10, and 15 in the day-6 to day-49 dataset with Louvain clustering at resolution 0.6. Cis-co-accessibility was determined over windows of 500kbp with 250kbp overlap. The accessibility score threshold for CCANs was set to 0.26. CCANs that overlapped with module gene bodies +1kb upstream were kept and associated to selected TFs by the presence of binding motifs in CCAN peaks associated with a gene body. All reported genomic regions refer to genome build hg38.

### Transfection

At day 19 of the differentiation protocol, cells were dissociated into a single cell suspension using TrypLE™ (Thermo Fisher Scientific) and plated under 2D culture conditions in CDM lacking TGF-β1, to allow them to attach and recover. On day 20, transfection was performed using Lipofectamine™ 3000 Transfection Reagent (Thermo Fisher Scientific; L3000001) according to the manufacturer’s instructions. Three experimental conditions were included: reference condition (untransfected control cells), empty vector using a pcDNA expression vector, and NFIA vector using a human NFIA (NM_001134673.4) cDNA ORF expression vector (Origene, Clone ID: OHu14494). To assess transfection efficiency, a parallel transfection was performed using a GFP-expressing pcDNA plasmid (Lonza). GFP expression was evaluated by fluorescence microscopy to estimate the proportion of transfected cells. On day 22 post-transfection, cells were dissociated. A fraction of the cells were pelleted into organoids by centrifugation for continued differentiatiation. Additional sampels were collected at day 25 post transfection for RNA isolation, while cultures were further maintained until day 35 in CDM, at which time were harvested for histological staining.

### RNA isolation and RT-qPCR

Total RNA content was isolated from cells on day-25, 5 days after transfection. Cells were lysed in 200 µl Trizol reagent (Thermo Fisher Scientific) and homogenized using micro pestles. RNA was extracted with chloroform and purified from the supernatant using the RNeasy Mini kit (Qiagen). RNA concentration was measured using a Nanodrop-1000 photospectrometer (Thermo Scientific). Synthesis of cDNA was performed with 150 ng of total RNA using a First Strand cDNA Synthesis kit according to manufacturer’s protocol (Roche Applied Science). The cDNA was diluted five times and expression levels of seven genes of interest were measured with QuantStudio 6 Real-Time PCR system using FastStart SYBR Green Master reaction mix (Roche Applied Science). Primer sequences are shown in **Supplementary Table 11**. Relative gene expression levels (-ΔCt) were calculated using the average of Ct values of Glyceraldehyde-3-Phosphate Dehydrogenase (*GAPDH*) and and Succinate dehydrogenase A (*SDHA*) as internal control (IC). ΔCt values of the gene of interest (GOI) were calculated as Ct_GOI_ – Ct_ic_. Relative expression levels (–ΔCt) were calculated using the mean of GAPDH and SDHA as reference genes. For each gene of interest (GOI), ΔCt was obtained as *Ct*₍GOI₎ – *Ct*₍IC₎.

### Neocartilage organoid size estimation

To evaluate neo-cartilage organoid, the size of constructs from each conditions was measured at the time of harvest. Samples were imaged using a brightfield microscope (Olympus CKX53), and two perpendicular diameters were recorded for each pellet using the arbitrary line tool in CellSens software (Olympus). The average of these diameters was used to calculate the cross-sectional area of each organoid, providing an estimate of organoid size.

### Histology

Neocartilage organoids at the indicated time point (day 35 and day 49) were fixed overnight in 4% formaldehyde and stored in 70% ethanol at 4°C. Samples were processed using a Tissue-Tek VIP 5 Jr. Processor (Sakura), embedded in paraffin, and sectioned at 5µm using a HistoCore Multicut microtome (Leica). Slides were dried, deparaffinized, and rehydrated prior to staining. Sections of 5µm thickness were obtained for all downstream analysis. Cartilage matrix composition was evaluated using 1% Alcian Blue 8-GX (Sigma-Aldrich) to visualize sulphated glycosaminoglycans (sGAGs), counterstained with Nuclear Fast Red (Sigma-Aldrich).

### Quantification and statistical analysis

All statistical analyses were performed in R. Clustering analysis was performed with Seurat WNN and Louvain based algorithm LSM. Differential expression analysis and differential activity analysis was performed on in psuedobulk with randomised pseudoreplicates in Delegate using edgeR testing and False Discovery Rate correction for multiple testing. Significance was determined with a 95% CI. Differential accessibility analysis used Bonferroni correction for multiple testing correction with a 95% CI. ClusterProfiler used standard enrichR procedures for pathway analysis using a 95% CI on *P* value and FDR to determine significance for terms.

## Acknowledgements

The Leiden University Medical Center (LUMC) supported this study and hiPSC cell line LUMC0004iCTRL10 was provided by the LUMC hiPSC core facility. This work was supported by the Dutch Research Council (NWO) [OCENW.GROOT.2019.079]. This work was done as part of the SCIMAP project and we would like to thank our collaborators from Twente University for their valuable input; Janine Post, Hil Meijer, and Lucas Klomp. We would like to thank the Leiden Genome Technology Center (LGTC) for their involvement. We would like to thank Fabienne de Jong for her assistance. Finally, we would like to thank the Dutch Society for Matrix Biology (NVMB) for their support.

## Data availability

Software and data used in this study will be made available for the full print release of this study. Data will be available on EGA. Software will be available on a Github repository via Zenodo.

## Brief overview of results and interest for the journal

In the current study we systematically mapped cell-fate trajectories from 7 time points during a 49-day chondrogenic hiPSC differentiation protocol using single-nucleus multimodal transcriptomic and chromatin accessibility profiling (scRNA-seq and scATAC-seq). Integrative analysis of dynamics and experimental validation revealed that *NFIA* and *NFIB* drove a firm chondrogenic split from a neurogenic trajectory.

We think that our study is interesting for the Cell Stem cells journal since it provides critical modifiable transcriptional regulators that can now be leveraged to implement regenerative cartilage therapies from hiPSCs.

## Figure legends

**Supplementary Figure 1:** pre-processing of single cell sequencing data, cell quality control. Cell status is indicated by colour and corresponds with **Fig. 1C**: High Quality (HQ) cells in red, High Quality in RNA (RNA) cells in green, High Quality in ATAC (ATAC) cells in blue, Low Quality (LQ) cells in blue. From top to bottom every row is a timepoint: day-0: I-V, day-3: VI-X, day-6: XI-XV, day-8: XVI-XX, day-14: XXI-XXV, day-21: XXVI-XXX, day-49: XXXI-XXXV. From left to right every column is a selection criterium: **RNA features** (I, VI, XI, XVI, XXI, XXVI, XXXI): Empty cells and doublets eliminated by selecting cells between the 15.8^th^ and 99^th^ percentile of features. **percentage mitochondrial RNA** (II, VII, XII, XVII, XXII, XXVII, XXXII): Cells with a percentage mitochondrial RNA over 95-99^th^ percentile eliminated. **Percentage ribosomal RNA** (III, VIII, XIII, XVIII, XXIII, XXVIII, XXXIII): Cells with a percentage ribosomal RNA over 85-90^th^ percentile were eliminated. **ATAC features** (IV, IX, XIV, XIX, XXIV, XXIX, XXXIV): Empty cells and doublets eliminated by selecting cells between the 22.8^th^ and 98.17^th^ percentile of peak region fragments. **ATAC Percentage peak fragments** (V, X, XV, XX, XXV, XXX, XXXV): Cells with a minority of fragments in peaks (<50%) eliminated.

**Supplementary Figure 2:** Dotplot of the top 10 most expressed gene per cluster in the whole dataset day-0 to day-49 with louvain clustering at resolution 0.2 (**Fig 2D**). **A**) Average scaled VST normalized expression for all cells in the cluster. **B**) Average VST normalized expression for all cells in the cluster

**Supplementary Fig. 3A** WNN UMAP projection of days 6-49 of differentiation with inferred trajectory in black, clustered to fit around the bifurcation point at day-21. Clusters are labeled and cell colour corresponds to labels. WNN UMAP projection of days 6-49 of differentiation with inferred trajectory in black, clustered to fit around the bifurcation point at day-21. Clusters are labeled and cell colour corresponds to labels.

**Supplementary Figure 3B:** Transcription factor activity assay of post-bifurcation neurogenic off-target cluster 8 as reference to post-bifurcation chondrogenic on-target cluster 15 as query.

**Supplementary Figure 3C:** Venn Diagram with for the differential activity between clusters surrounding the bifurcation at day 21(**Fig. 5A-B**). Comparisons included are pre-bifurcation cluster 10 as reference vs post-bifurcation on-target cluster 15 as query, pre-bifurcation cluster 10 as reference to post-bifurcation off-target cluster 8 as query, and post-bifurcation off-target cluster 8 as reference vs post-bifurcation on-target cluster 15 as query. Each comparison is stratified by consensus of log fold change in differential expression and differential motif enrichment as indicated in each supplementary table. Specificity is defined as the exclusivity positive or negative fold change to the on- or off-target direction from pre- to post-bifurcation as modulated by the differential expression between post-bifurcation clusters.

**Supplementary Figure 4:** Nuclear Factor one centric transcription factor-target network for the on-target direction trajectory-1. This network is a visual representation of **Supplementary Table 9**. Central are the NFI genes NFIA, NFIB, and NFIX their differentially expressed module gene targets, co-factors for their differentially expressed module gene targets, and transcription factors which target NFI genes. All transcription factors considered were found to be “specifically upregulated in on-target” or “specifically upregulated in on-target and downregulated in off-target” (**Supplementary Figure 3**). Moran’s I was taken from **Supplementary table 1** and log fold change expression from differential expression analysis between pre-bifurcation cluster 10 as reference vs post-bifurcation on-target cluster 15 as query (**Supplementary Table 6**).

**Supplementary Figure 5:** Coverage plot of cis-co-accessibility network nr. 520 which covers the NFIA gene locus. This CCAN covers genomic range chr1:60547451-61589714 in hg38. From top to bottom, left to right: The first three tracks show normalized signal (fragment count) across the range of interest for the pre-bifurcation cluster 10, post-bifurcation chondrogenic cluster 15, post-bifurcation neurogenic cluster 8 (Fig. 5B). On the right side are violin plots that show the VST normalized expression for NFIA per corresponding cluster. The fourth track shows the forward and reverse gene loci in the genomic range of interest. The fifth track shows peaks called from fragments. The sixth track shows transcription factor binding sites in peaks that make up CCAN nr. 520. Only TFs that target NFIA are included for the chondrogenic trajectory (FOXA1, FOXP2, NFATC2, RUNX2) and the neurogenic trajectory (NHLH1, ONECUT1, ONECUT3, ZEB1). The seventh track shows the links between peaks that makes up CCAN nr. 520 colored by co-accessibility score.

**Supplementary Figure 6:** NFIA transfection experiment. **A)** Diagram of the experimental progress. **B)** Brightfield microscopy images at 4x magnification (500 µm black bar) of neocartilage organoids at day 35 in three conditions: NFIA+ in duplo, empty vector in triplo, and wild type in triplo.

