## Supplementary Figures for "Nuclear Factor I genes drive chondrogenic cell-fate commitment"

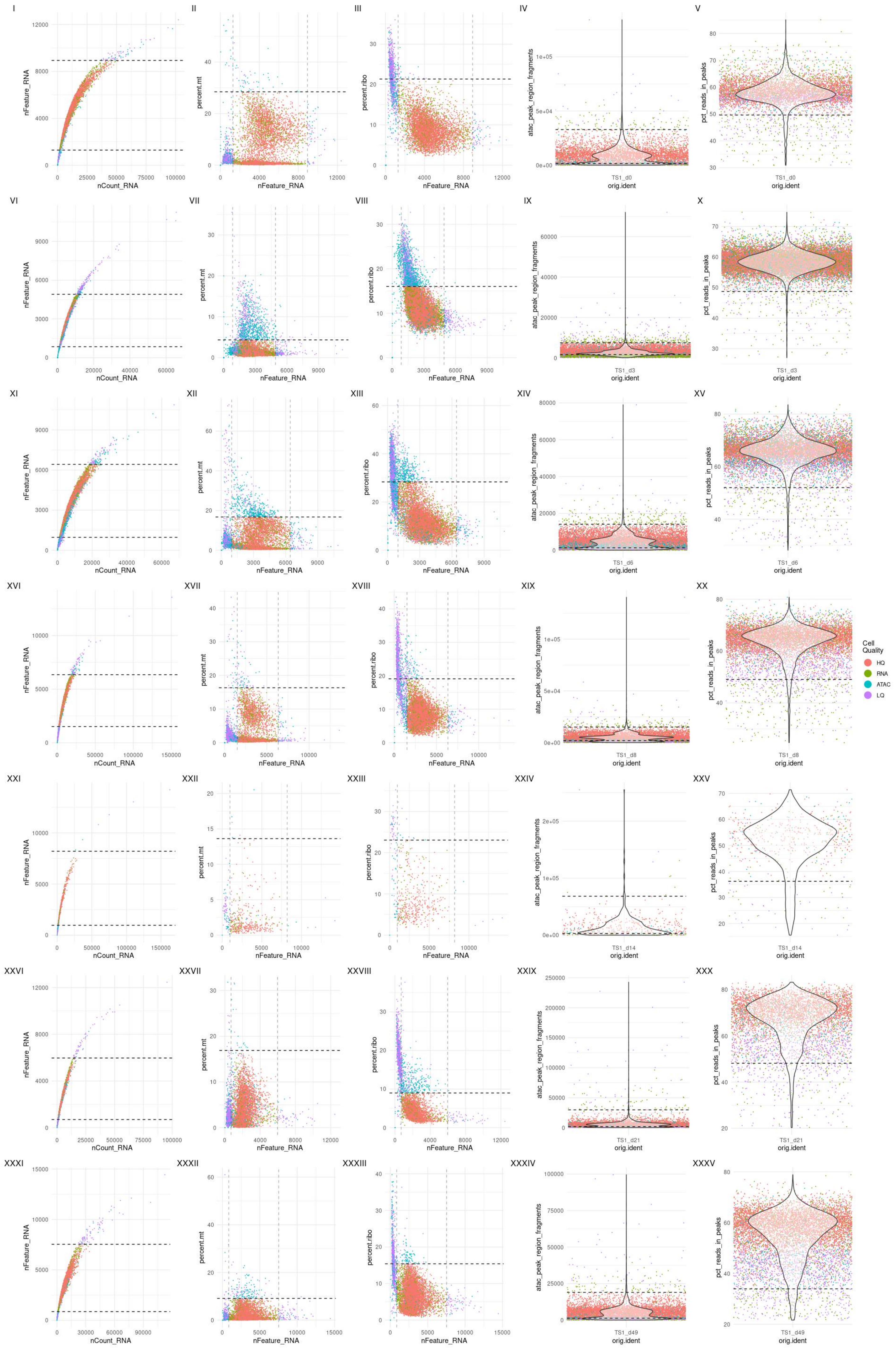
I
II
III
IV
V
VI
VII
VIII
IX
X
XI
XII
XIII
XIV
XV
XVI
XVII
XVIII
XIX
XX
XXI
XXII
XXIII
XXIV
XXV
XXVI
XXVII
XXVIII
XXIX
XXX
XXXI
XXXII
XXXIII
XXXIV
XXXV
nFeature_RNA
nCount_RNA
percent.mt
percent.ribo
atac_peak_region_fragments
pct_reads_in_peaks
orig.ident
TS1_d0
TS1_d3
TS1_d6
TS1_d8
TS1_d14
TS1_d21
TS1_d49
Cell Quality
HQ
RNA
ATAC
LQ

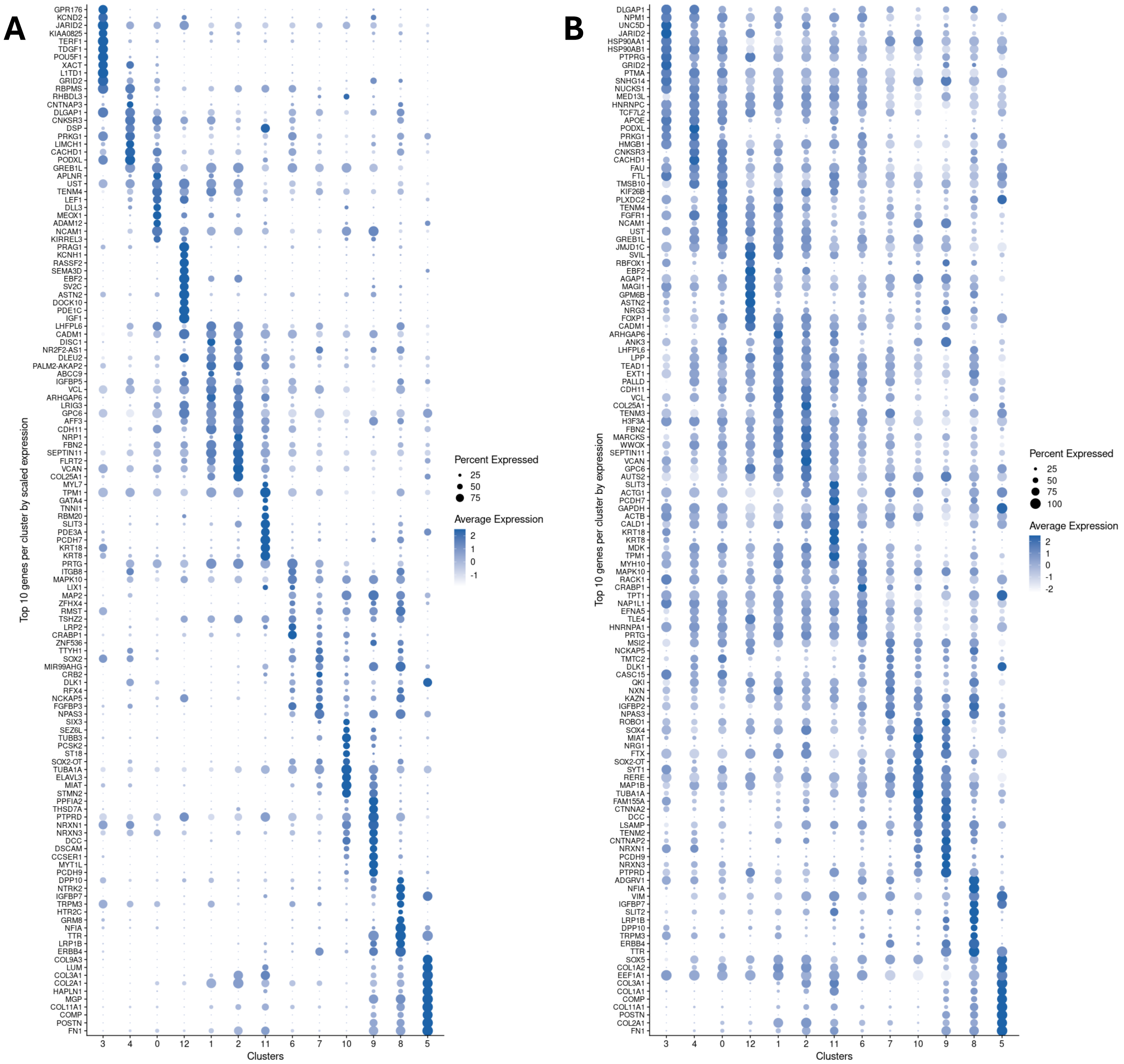

A
GPR176
KCND2
JARID2
KIAA0825
TERF1
TDGF1
POU5F1
XACT
L1TD1
GRID2
RBPMS
RHBDL3
CNTNAP3
DLGAP1
CNKSR3
DSP
PRKG1
LIMCH1
CACHD1
PODXL
GREB1L
APLNR
UST
TENM4
LEF1
DLL3
MEOX1
ADAM12
NCAM1
KIRREL3
PRAG1
KCNH1
RASSF2
SEMA3D
EBF2
SV2C
ASTN2
DOCK10
PDE1C
IGF1
LHFPL6
CADM1
DISC1
NR2F2-AS1
DLEU2
PALM2-AKAP2
ABCC9
IGFBP5
VCL
ARHGAP6
LRIG3
GPC6
AFF3
CDH11
NRP1
FBN2
SEPTIN11
FLRT2
VCAN
COL25A1
MYL7
TPM1
GATA4
TNNI1
RBM20
SLIT3
PDE3A
PCDH7
KRT18
KRT8
PRTG
ITGB8
MAPK10
LIX1
MAP2
ZFHX4
RMST
TSHZ2
LRP2
CRABP1
ZNF536
TTYH1
SOX2
MIR99AHG
CRB2
DLK1
RFX4
NCKAP5
FGFBP3
NPAS3
SIX3
SEZ6L
TUBB3
PCSK2
ST18
SOX2-OT
TUBA1A
ELAVL3
MIAT
STMN2
PPFIA2
THSD7A
PTPRD
NRXN1
NRXN3
DCC
DSCAM
CCSER1
MYT1L
PCDH9
DPP10
NTRK2
IGFBP7
TRPM3
HTR2C
GRM8
NFIA
TTR
LRP1B
ERBB4
COL9A3
LUM
COL3A1
COL2A1
HAPLN1
MGP
COL11A1
COMP
POSTN
FN1
Top 10 genes per cluster by scaled expression
3 4 0 12 1 2 11 6 7 10 9 8 5
Clusters
Percent Expressed
25
50
75
Average Expression
2
1
0
-1
B
DLGAP1
NPM1
UNC5D
JARID2
HSP90AA1
HSP90AB1
PTPRG
GRID2
PTMA
SNHG14
NUCKS1
MED13L
HNRNPC
TCF7L2
APOE
PODXL
PRKG1
HMGB1
CNKSR3
CACHD1
FAU
FTL
TMSB10
KIF26B
PLXDC2
TENM4
FGFR1
NCAM1
UST
GREB1L
JMJD1C
SVIL
RBFOX1
EBF2
AGAP1
MAGI1
GPM6B
ASTN2
NRG3
FOXP1
CADM1
ARHGAP6
ANK3
LHFPL6
LPP
TEAD1
EXT1
PALLD
CDH11
VCL
COL25A1
TENM3
H3F3A
FBN2
MARCKS
WWOX
SEPTIN11
VCAN
GPC6
AUTS2
SLIT3
ACTG1
PCDH7
GAPDH
ACTB
CALD1
KRT18
KRT8
MDK
TPM1
MYH10
MAPK10
RACK1
CRABP1
TPT1
NAP1L1
EFNA5
TLE4
HNRNPA1
PRTG
MSI2
NCKAP5
TMTC2
DLK1
CASC15
QKI
NXN
KAZN
IGFBP2
NPAS3
ROBO1
SOX4
MIAT
NRG1
FTX
SOX2-OT
SYT1
RERE
MAP1B
TUBA1A
FAM155A
CTNNA2
DCC
LSAMP
TENM2
CNTNAP2
NRXN1
PCDH9
NRXN3
PTPRD
ADGRV1
NFIA
VIM
IGFBP7
SLIT2
LRP1B
DPP10
TRPM3
ERBB4
TTR
SOX5
COL1A2
EEF1A1
COL3A1
COL1A1
COMP
COL11A1
POSTN
COL2A1
FN1
Top 10 genes per cluster by expression
3 4 0 12 1 2 11 6 7 10 9 8 5
Clusters
Percent Expressed
25
50
75
100
Average Expression
2
1
0
-1
-2

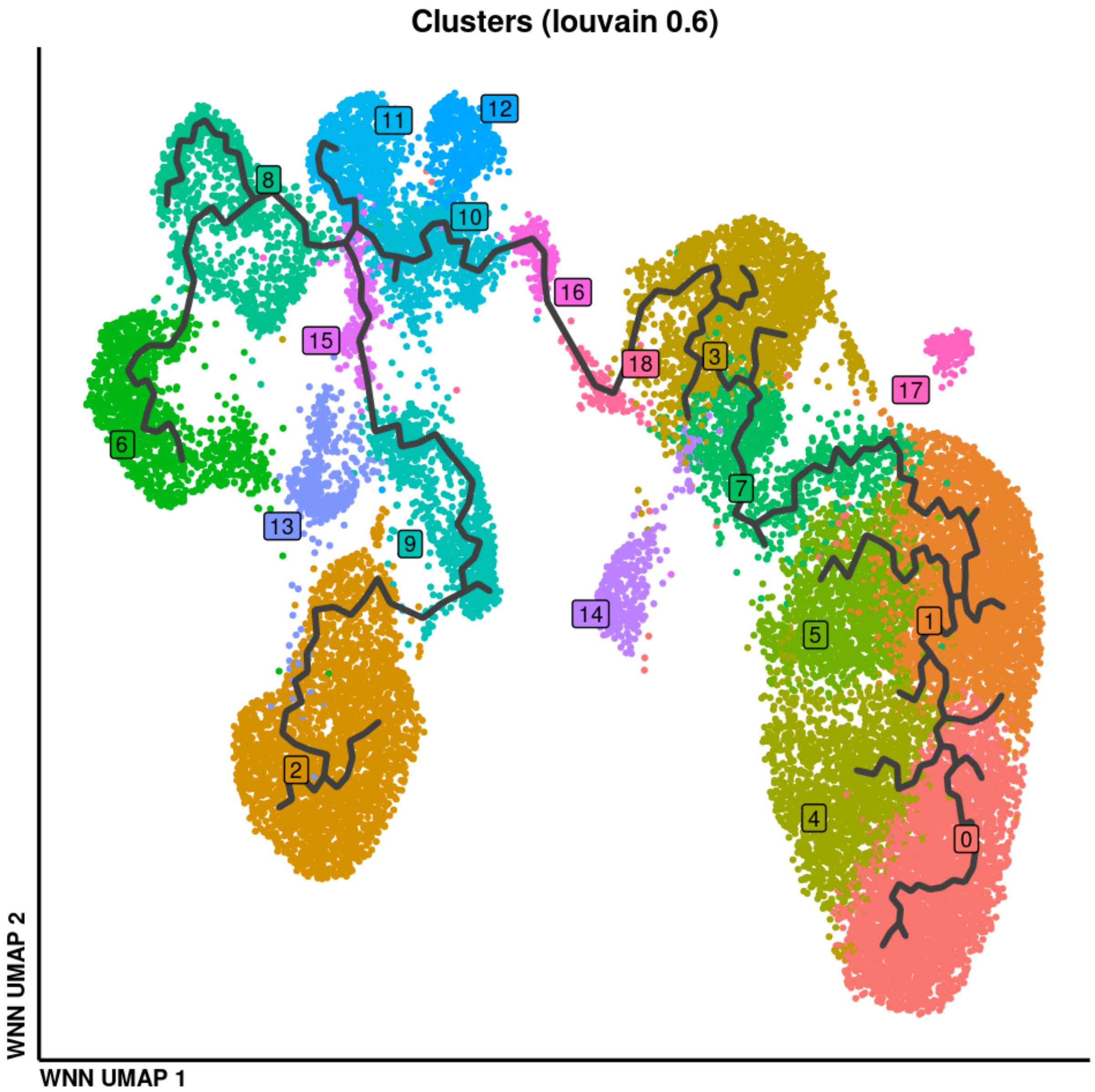
Clusters (louvain 0.6)
11
12
8
10
16
15
18
3
17
6
7
13
9
14
5
1
2
4
0
WNN UMAP 2
WNN UMAP 1

### TF activity reference post-bifurcation off-target (8) to query post-bifurcation on-target (15)

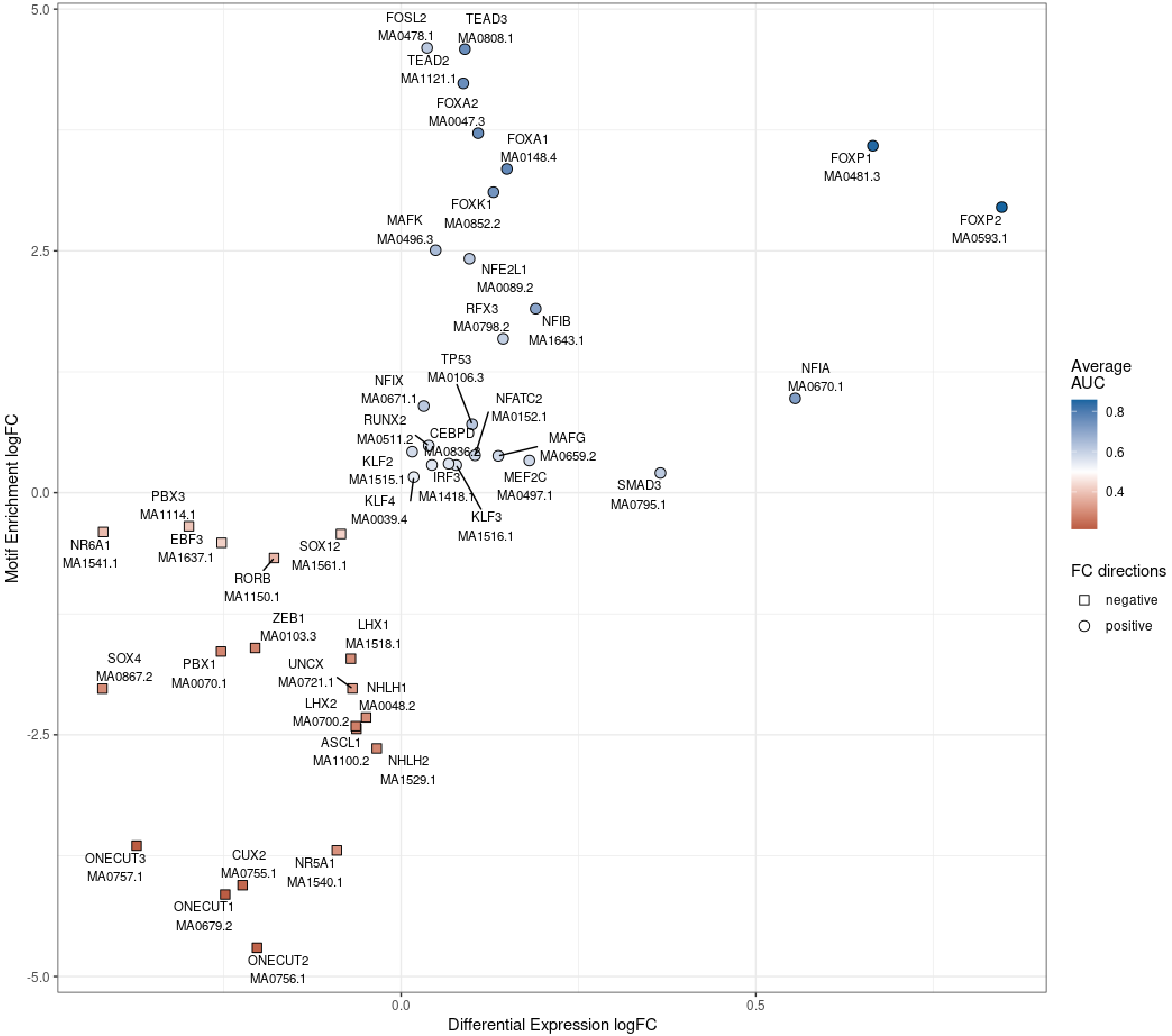

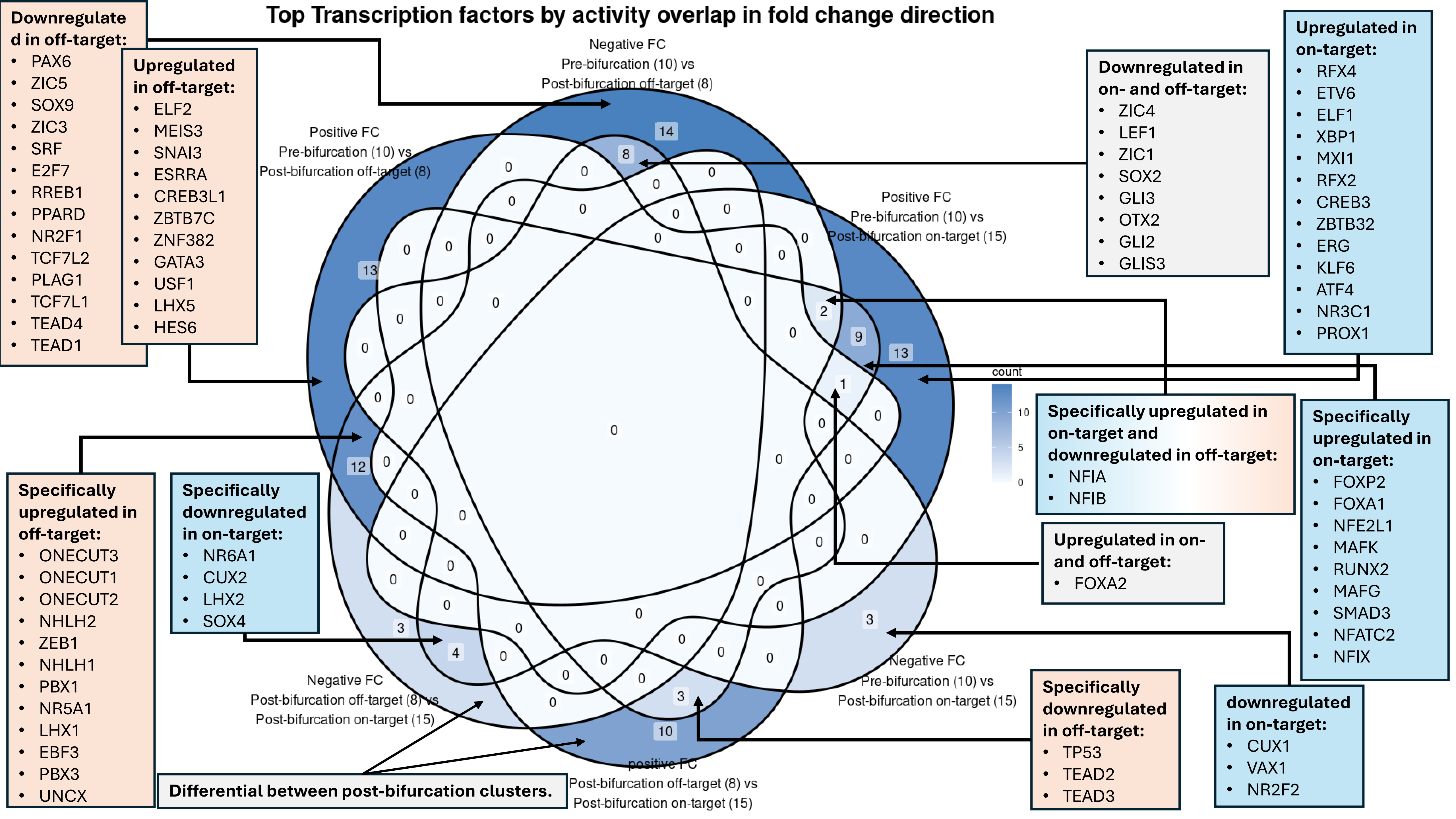
Top Transcription factors by activity overlap in fold change direction
Negative FC
Pre-bifurcation (10) vs
Post-bifurcation off-target (8)
Positive FC
Pre-bifurcation (10) vs
Post-bifurcation off-target (8)
Positive FC
Pre-bifurcation (10) vs
Post-bifurcation on-target (15)
Negative FC
Pre-bifurcation (10) vs
Post-bifurcation on-target (15)
Negative FC
Post-bifurcation off-target (8) vs
Post-bifurcation on-target (15)
positive FC
Post-bifurcation off-target (8) vs
Post-bifurcation on-target (15)
14
8
13
12
2
9
13
1
3
4
3
3
10
count
10
5
0
Downregulated in off-target: PAX6, ZIC5, SOX9, ZIC3, SRF, E2F7, RREB1, PPARD, NR2F1, TCF7L2, PLAG1, TCF7L1, TEAD4, TEAD1
Upregulated in off-target: ELF2, MEIS3, SNAI3, ESRRA, CREB3L1, ZBTB7C, ZNF382, GATA3, USF1, LHX5, HES6
Downregulated in on- and off-target: ZIC4, LEF1, ZIC1, SOX2, GLI3, OTX2, GLI2, GLIS3
Upregulated in on-target: RFX4, ETV6, ELF1, XBP1, MXI1, RFX2, CREB3, ZBTB32, ERG, KLF6, ATF4, NR3C1, PROX1
Specifically upregulated in on-target and downregulated in off-target: NFIA, NFIB
Specifically upregulated in on-target: FOXP2, FOXA1, NFE2L1, MAFK, RUNX2, MAFG, SMAD3, NFATC2, NFIX
Upregulated in on- and off-target: FOXA2
Specifically upregulated in off-target: ONECUT3, ONECUT1, ONECUT2, NHLH2, ZEB1, NHLH1, PBX1, NR5A1, LHX1, EBF3, PBX3, UNCX
Specifically downregulated in on-target: NR6A1, CUX2, LHX2, SOX4
Specifically downregulated in off-target: TP53, TEAD2, TEAD3
downregulated in on-target: CUX1, VAX1, NR2F2
Differential between post-bifurcation clusters.

NFI Transcription Factor - Target network: on-target

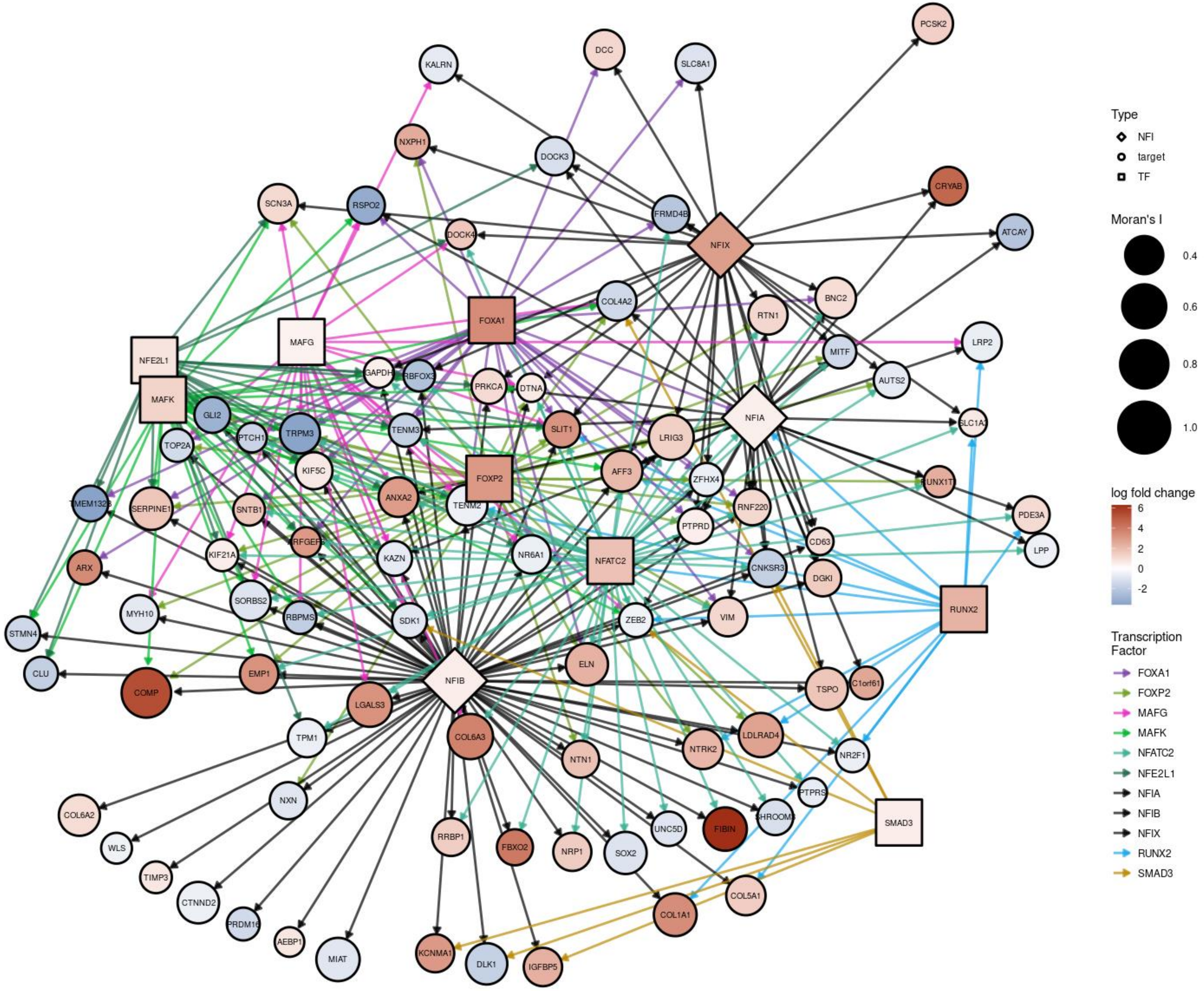

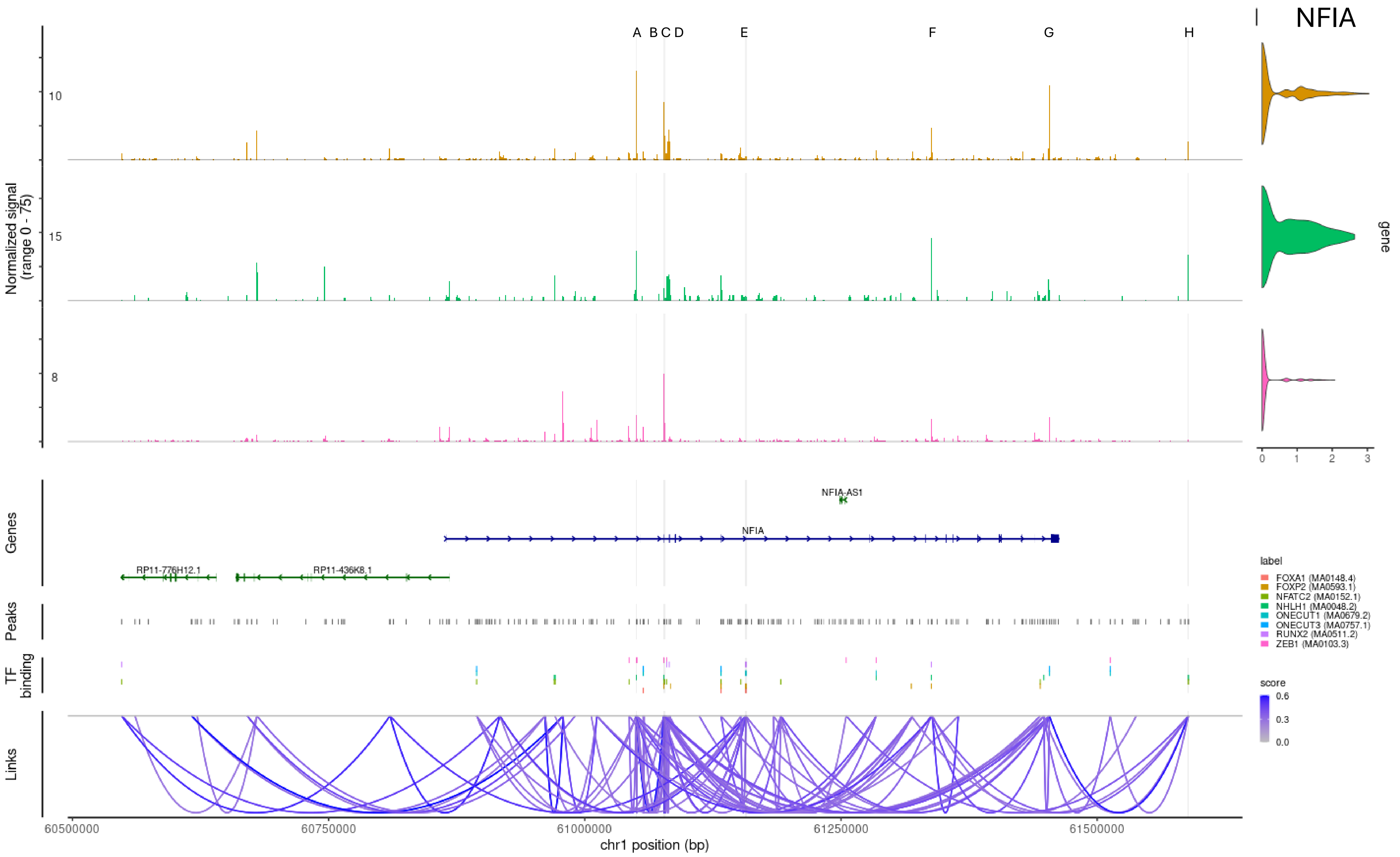
NFIA
A B C D
E
F
G
H
Normalized signal
(range 0 - 75)
10
15
8
gene
0
1
2
3
Genes
NFIA-AS1
NFIA
RP11-776H12.1
RP11-436K8.1
Peaks
TF binding
Links
label
FOXA1 (MA0148.4)
FOXP2 (MA0593.1)
NFATC2 (MA0152.1)
NHLH1 (MA0048.2)
ONECUT1 (MA0679.2)
ONECUT3 (MA0757.1)
RUNX2 (MA0511.2)
ZEB1 (MA0103.3)
score
0.6
0.3
0.0
60500000
60750000
61000000
61250000
61500000
chr1 position (bp)

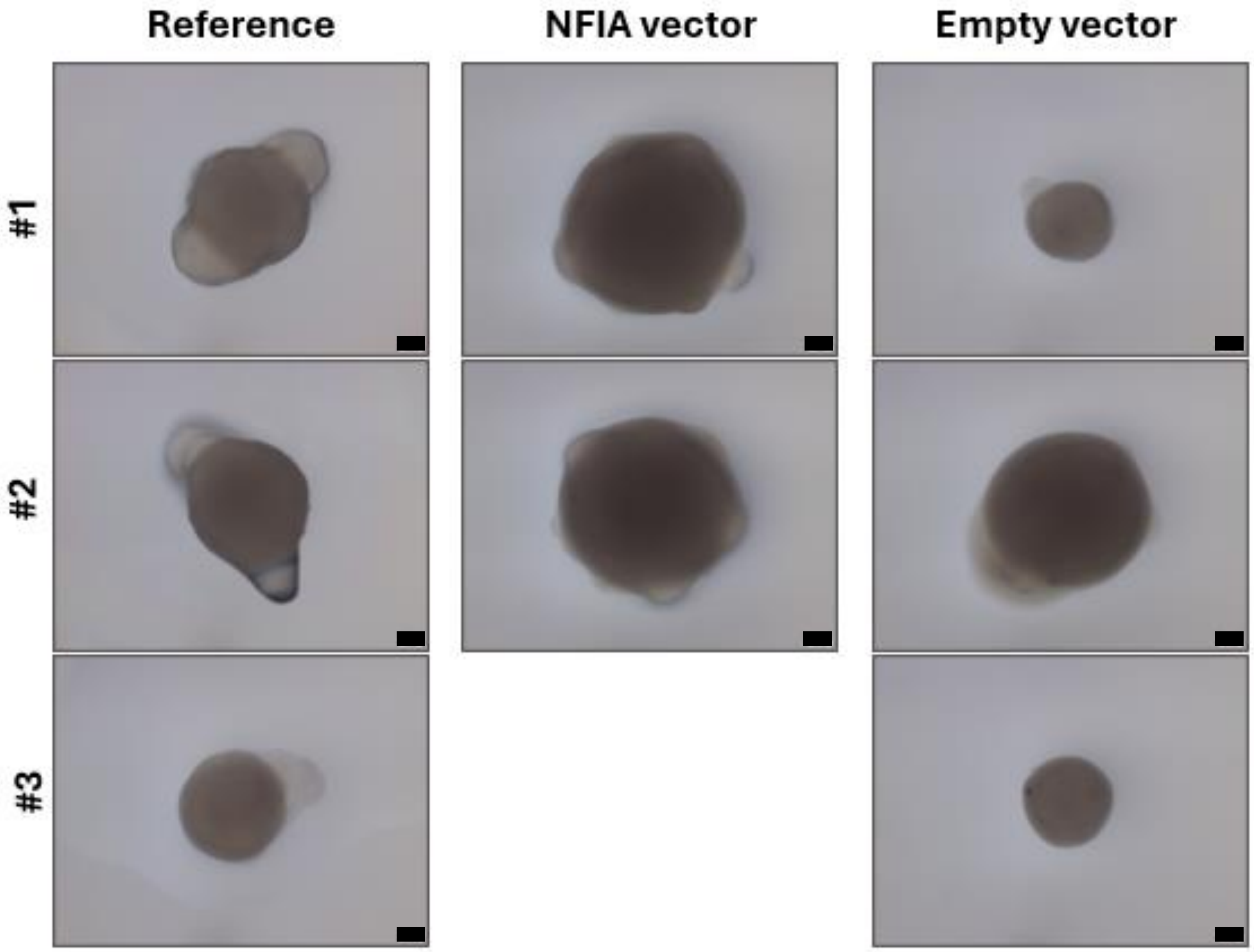
Reference
NFIA vector
Empty vector
#1
#2
#3
